# Regenerative neurogenic response from glia requires insulin driven neuron-glia communication

**DOI:** 10.1101/721498

**Authors:** Neale Harrison, Elizabeth Connolly, Alicia Gascón Gubieda, Zidan Yang, Benjamin Altenhein, Maria Losada-Perez, Marta Moreira, Alicia Hidalgo

**Author notes:** These authors contributed equally (Neale Harrison, Elizabeth Connolly). Author for correspondence: Professor Alicia Hidalgo ph 00 44 (0)121 4145416.

## Abstract

Some animals can regenerate their central nervous system (CNS) after injury by inducing de novo neurogenesis: discovering the underlying mechanisms would help promote regeneration in the damaged human CNS. Glial cells could be the source of regenerative neurogenesis, but this is debated. The glia transmembrane protein Neuron-Glia antigen-2 (NG2) may have a key role in sensing injury-induced neuronal signals, however these have not been identified. Here, we used *Drosophila* genetics to search for functional neuronal partners of the *NG2* homologue *kon-tiki (kon),* and identified Islet Antigen-2 (Ia-2), required in neurons for insulin secretion. Alterations in Ia-2 function induced neural stem cell fate, injury increased *ia-2* expression and induced ectopic neural stem cells. Using genetic epistasis analysis and lineage tracing, we demonstrate that Ia-2 functions with Kon to regulate *Drosophila* insulin-like peptide 6 (Dilp-6) which in turn generates both more glial cells and neural stem cells from glia. Ectopic neural stem cells can divide, and limited de novo neurogenesis could be traced back to glial cells. Altogether, Ia-2 and Dilp-6 drive a neuron-glia relay that restores glia, and reprograms glia into neural stem cells for CNS regeneration.

## INTRODUCTION

The central nervous system (CNS) can regenerate after injury in some animals, and this involves de novo neurogenesis (Tanaka and Ferretti 2009). Newly formed neurons integrate into functional neural circuits, enabling the recovery of function and behavior, which is how CNS regeneration is measured (Tanaka and Ferretti 2009). The human CNS does not regenerate after injury. However, in principle it could, as we continue to produce new neurons throughout life, that integrate into functional circuits (Tanaka and Ferretti 2009; Gage 2019). Through understanding the molecular mechanisms underlying natural regenerative neurogenesis in animals, we might be able to provoke de novo neurogenesis in the human CNS, to promote regeneration after damage or neurodegenerative diseases. Regenerative neurogenesis across animals may reflect an ancestral, evolutionarily conserved genetic mechanism, which manifests itself to various degrees in regenerating and non-regenerating animals (Tanaka and Ferretti 2009). Accordingly, it may be possible to discover molecular mechanisms of injury-induced neurogenesis in the fruit-fly *Drosophila,* which is a powerful genetic model organism.

Regenerative neurogenesis could occur through activation of quiescent neural stem cells, de-differentiation of neurons or glia, or direct conversion of glia to neurons (Tanaka and Ferretti 2009; Falk and Gotz 2017). Across many regenerating animals, new neurons originate mostly from glial cells (Tanaka and Ferretti 2009; Falk and Gotz 2017). In the mammalian CNS, radial glial cells behave like neural stem cells to produce neurons during development. Remarkably, whereas NG2-glia (also known as oligodendrocyte progenitor cells, OPCs) produce only glia (oligodendrocytes and astrocytes) in development, they can also produce neurons in the adult and upon injury (Dimou and Gotz 2014; Falk and Gotz 2017; Valny et al. 2017; Du et al. 2020) – although this remains controversial. Discovering the molecular mechanisms of a neurogenic response of glia is of paramount urgency.

NG2-glia are progenitor cells in the adult human brain, constituting 5-10% of cells in the total CNS, and remain proliferative throughout life (Dimou and Gotz 2014). In development, NG2-glia are progenitors of astrocytes, OPCs and oligodendrocytes, but postnatally and upon injury they can also produce neurons (Dimou and Gotz 2014; Torper et al. 2015; Falk and Gotz 2017; Valny et al. 2017; Du et al. 2020). They can also be directly reprogrammed into neurons that integrate into functional circuits (Torper et al. 2015; Pereira et al. 2017). The diversity and functions of NG2-glia are not yet fully understood, but they are particularly close to neurons: they receive and respond to action potentials integrating them into calcium signals, monitor and modulate the state of neural circuits by regulating channels, secrete chondroitin sulfate proteoglycan perineural nets, and they also induce their own proliferation to generate more NG2-glia, astrocytes that sustain neuronal physiology, and oligodendrocytes that enwrap axons (Dimou and Gotz 2014; Sakry and Trotter 2016; Sun et al. 2016; Du et al. 2020). NG-2 glia have key roles in brain plasticity, homeostasis and repair in close interaction with neurons (Dimou and Gotz 2014; Sakry and Trotter 2016; Du et al. 2020), but to what extent this depends on the *NG2* gene and protein, is not known.

*NG2 (also known as chondroitin sulfate proteoglycan 4, CSPG4)* is expressed by NG2-glia and pericytes, but not by oligodendrocytes, neurons, or astrocytes (Cahoy et al 2008). NG2 is a transmembrane protein that can be cleaved upon neuronal stimulation to release a large secreted extracellular domain and an intracellular domain (Sakry et al. 2014; Sakry and Trotter 2016). The intracellular domain (ICD, NG2^ICD^) is mostly cytoplasmic, and it induces protein translation and cell cycle progression (Nayak et al. 2018). NG2^ICD^ lacks a DNA binding domain and therefore does not function as a transcription factor, but it has a nuclear WW4 domain and nuclear localization signals, and can regulate gene expression (Sakry et al. 2015; Sakry and Trotter 2016; Nayak et al. 2018). It is thought that NG2 functions as a receptor, triggering nuclear signalling in response to ligands/partners (Sakry et al. 2014; Sakry and Trotter 2016). NG2 protein is abundant in proliferating NG2-glia and glioma (Sakry et al. 2015; Sakry and Trotter 2016; Nayak et al. 2018), and it is required for OPC proliferation and migration in development and in response to injury (Kucharova and Stallcup 2010; Kucharova et al. 2011; Biname et al. 2013). Given the close relationship of NG2-glia with neurons, it is anticipated that key partners of NG2 are produced from neurons, but these remain largely unknown.

The fruit-fly *Drosophila* is particularly powerful for discovering novel molecular mechanisms. The *Drosophila NG2* homologue is called *kon-tiki (kon)* or *perdido* (Estrada et al. 2007; Schnorrer et al. 2007; Perez-Moreno et al. 2017). Kon functions in glia, promotes glial proliferation and glial cell fate determination in development and upon injury, and promotes glial regeneration and CNS injury repair (Losada-Perez et al. 2016). Kon works in concert with the receptor Notch and the transcription factor Prospero (Pros) to drive the glial regenerative response to CNS injury (Kato et al. 2011; Losada-Perez et al. 2016). It is normally found in low levels in the larval CNS, but injury induces a Notch-dependent increase in *kon* expression in glia (Losada-Perez et al. 2016). Together, Notch signaling and Kon induce glial proliferation; Kon also initiates neuropile glial differentiation and *pros* expression, and Pros maintains glial cell differentiation (Griffiths and Hidalgo 2004; Kato et al. 2011; Losada-Perez et al. 2016). This glial regenerative response to injury is homeostatic and time-limited, as two negative feedback loops halt it: Kon represses Notch, and Pros represses *kon* expression, preventing further cell division (Kato et al. 2011; Losada-Perez et al. 2016). The relationship between these genes is also conserved in the mouse, where the homologue of *pros, Prox1,* is expressed together with *Notch1* in NG2-glia, and following cell division, it represses NG2-glia proliferation and promotes oligodendrocyte differentiation (Kato et al. 2015). Together, Notch, Kon and Pros form a homeostatic gene network that sustains neuropile glial integrity throughout life and drives glial regeneration upon injury (Hidalgo and Logan 2017; Kato et al. 2018). As Kon is up-regulated upon injury and provokes glial proliferation and differentiation, it is the key driver of the glial regenerative response to CNS injury.

A critical missing link to understand CNS regeneration was the identification of neuronal partners of glial NG2/Kon that could induce regenerative neurogenesis. We had observed that injury to the *Drosophila* larval CNS also resulted in spontaneous, yet incomplete, repair of the axonal neuropile (Kato et al. 2011). This strongly suggested that injury might also induce neuronal events, such as axonal regrowth, or generation of new neurons. Thus, we asked whether Kon may interact with neuronal factors that could contribute to regenerative neurogenesis after injury. Here, we report that relay of insulin signalling involving neuronal Ia-2 and glial Kon, drives in vivo reprogramming of neuropile glia into neural stem cells.

## RESULTS

### Ia-2 is a functional partner of Kon produced in neurons

To search for functional neuronal partners of Kon, we carried out genetic screens that aimed to identify genes expressed in neurons that had non-autonomous effects on glia. We exploited the fact that over-expression of *kon* elongates the larval ventral nerve cord (VNC) (Losada-Perez et al. 2016), and tested whether RNAi knock-down of candidate genes in neurons or glia could rescue this phenotype (Figure 1 – Figure Supplements 1 and 2). To validate the approach, we first tested genes predicted or known to interact with *kon* and/or *NG2* (Schnorrer et al. 2007; Perez-Moreno et al. 2017). Indeed, knock-down of known interactors, such as *integrins* (Perez-Moreno et al. 2017), factors involved in Notch signaling (e.g. *Mtm, Akap200*), secretases (i.e. *kuz, kuz-l*) that cleave both Notch and NG2/Kon (Sakry and Trotter 2016), and phosphatases *Prl1* and *Ptp99A* (Song et al. 2012), all rescued the phenotype, validating the approach (Figure 1 – Figure Supplements 1 and 2). We tested knock-down of other genes encoding phosphatases and transmembrane proteins expressed in neurons. Knocking-down phosphatases *ptp99A, ptp69D* and *ptp4E* from neurons rescued the phenotype, but most prominent was knock-down of phosphatase *lar,* a negative regulator of insulin signalling (Figure 1 – Figure Supplement 2A-D). Notably, knock-down of other insulin related factors including *Akt* and *ia-2,* also caused some rescue (Figure 1 – Figure Supplement 2A-D). However, multiple genes can affect VNC length, and these rescue phenotypes may not necessarily reflect specific gene interactions. Thus, we next asked whether altering *kon* function affected the expression of a group of genes selected from the above screens. Kon can influence gene expression, as *kon* mutations cause loss of glial gene expression (Losada-Perez et al. 2016). Using quantitative real-time reverse transcription PCR (qRT-PCR) on dissected larval CNSs, we found that *kon* knock-down in neurons (with *kon^c452^, elavGAL4>UAS-konRNAi*) or glia (with *kon^c452^, repoGAL4>UAS-konRNAi*) had no effect on the expression of most phosphatases, including *lar*, or other tested genes, but by contrast, resulted in an approximately 3-fold increase in *ia-2* mRNA levels (Figure 1 – Figure Supplement 3A). Conversely, over-expression of full-length *kon* in either neurons or glia down-regulated *ia-2* mRNA levels by 25% (Figure 1 – Figure Supplement 3B). We validated these results by increasing the repeats of the most promising subset of genes (Figure 1 – Figure Supplement 3C,D), and this confirmed the strongest effect of *kon* loss and gain of function on *ia-2* (Figure 1A). Accordingly, Kon function in glia prevents *ia-2* expression. Next, we asked whether knock-down or over-expression of *ia-2* in neurons (with *elavGAL4*) had any effect on *kon* mRNA levels, but none did (Figure 1B). However, over-expression of *ia-2* in glia (with *repoGAL4>ia-2[GS11438]*) decreased *kon* mRNA levels (Figure 1B). As Kon functions in glia (Losada-Perez et al. 2016), these data indicated that *kon* and *ia-2* restrict each other’s expression to glia or neurons, respectively, and/or that Ia-2 is restricted to neurons. Either way, these data showed that *ia-2* and *kon* interact genetically.

**FIGURE 1.**
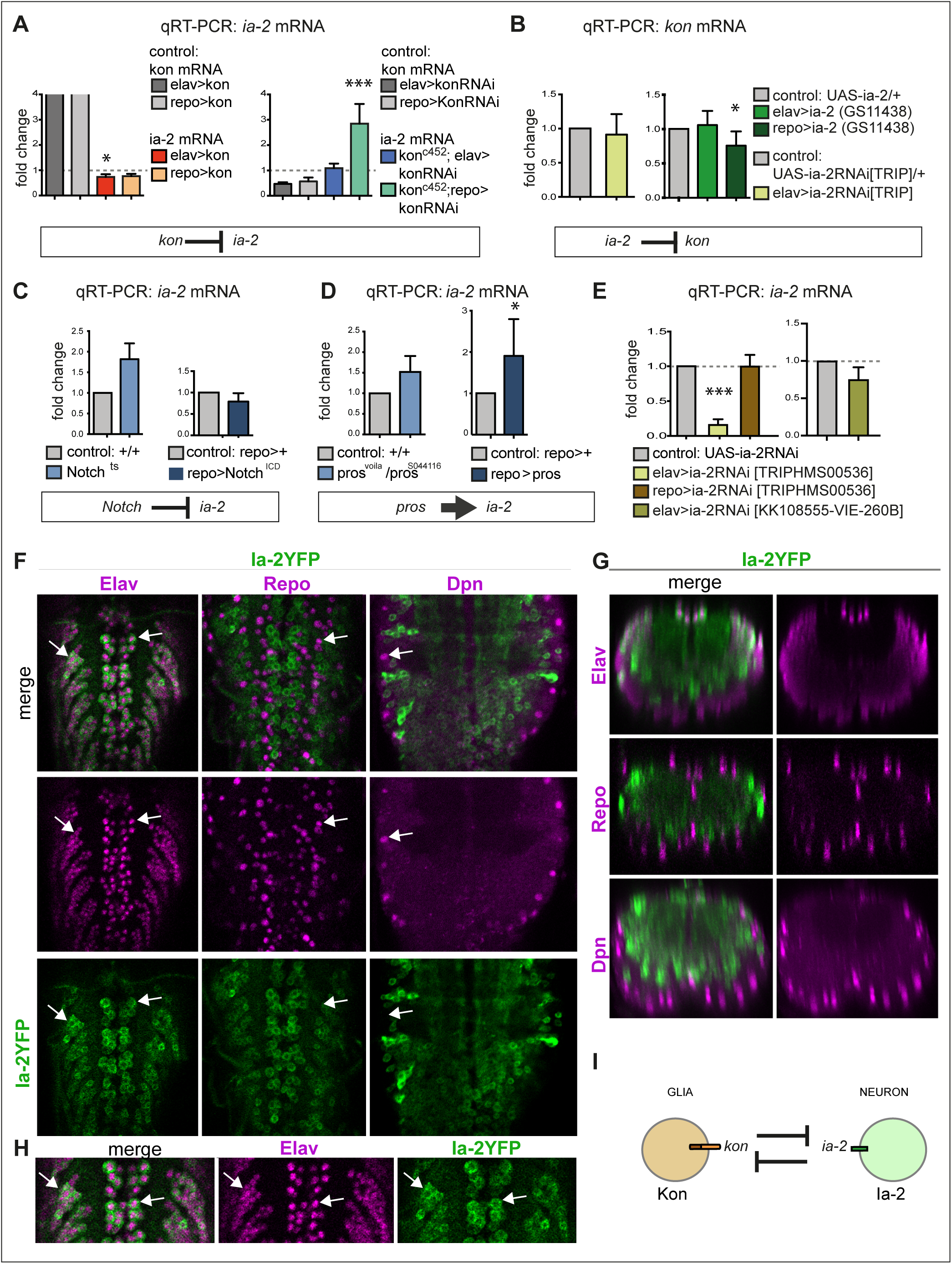
*ia-2* interacts genetically with *kon, Notch* and *pros.* **(A)** Quantitative real-time PCR (qRT-PCR) showing that gain of *kon* function reduced *ia-2* mRNA levels by 25% (One Way ANOVA p=0.045), whereas loss of *kon* function in glia caused practically a 3-fold increase in *ia-2* mRNA levels (*genotype: kon^c452^/UASkonRNAi; repoGAL4/+*; One Way ANOVA p<0.0001). Post-hoc Dunnett’s test multiple comparisons to control. N=4 replicates. **(B)** qRT-PCR showing that over-expression of *ia-2* in glia downregulated *kon* mRNA levels. Left: Unpaired Student t-test with Welch correction p=0.457. Right: One Way ANOVA p<0.045, post-hoc Dunnett’s test multiple comparisons to control. N=4-6 replicates. **(C)** Ia-2 is functionally related to Notch: qRT-PCR showing that *ia-2* mRNA levels increased in *N^ts^* mutant larvae at the restrictive temperature of 25°C. Unpaired Student t-test with Welch correction Left: p=0.4123; Right: p=0.2182. N=3 replicates. **(D)** *ia-2* is functionally related to *pros*: qRT-PCR showing that over-expression of *pros* in glia increased *ia-2* mRNA levels by 2-fold. Unpaired Student t-test with Welch correction. Left: p=0.1368; Right: p=0.0428. N=3 replicates. **(E)** qRT-PCR showing that *UAS-ia-2* RNAi[TRIPHMS00536] knock-down in neurons (with *elavGAL4*) lowered *ia-2* mRNA levels to 20%, whereas in glia it has no effect, meaning that *ia-2* is expressed in neurons. A second *UAS-ia-2RNAi[KK108555-VIE-260B]* line lowered mRNA levels by 25%. One Way ANOVA p=0.0004, post-hoc multiple comparisons to control Dunnett’s test. N=3 replicates. **(F,G,H)** Fusion protein Ia-2YFP revealed expression exclusively in neurons, as all Ia-2YFP+ cells were also Elav+, but Repo^—^ and Dpn^—^. Genotype: ia-2[CPTI100013]. N=4-16 larval VNCs. **(I)** Illustration showing that *kon* and *ia-2* functions are restricted to glia and neurons, respectively, and they mutually exclude each other. **(G)** transverse views; **(F,H)** horizontal views; **(H)** higher magnification views). With more than two sample types, asterisks indicate multiple comparison post-hoc tests to controls: *p<0.05, **p<0.01, ***p<0.001, ****p<0.0001. For full genotypes and further statistical analysis details see Table Supplement 1.

Our genetic and qRT-PCR based screens had identified genetic interactions between *kon* and *lar, Akt* and *ia-2.* LAR is involved in neuronal axon guidance, and is responsible for de-phosphorylating, and thus inactivating, insulin receptor signaling (Mooney et al. 1997; Wills et al. 1999). Akt is a key effector of insulin receptor signalling downstream (van der Heide et al. 2006). Ia-2 is a highly evolutionarily conserved phosphatase-dead transmembrane protein phosphatase required in dense core vesicles for the secretion of insulin, insulin-related factor-1 (IGF-1) and neurotransmitters; it also has synaptic functions and influences behaviour and learning (Cai et al. 2001; Harashima et al. 2005; Hu et al. 2005; Henquin et al. 2008; Cai et al. 2009; Nishimura et al. 2010; Cai et al. 2011; Carmona et al. 2014). Rather unexpectedly, our findings suggested that Kon is involved in insulin signalling.

To ask whether and how Ia-2 might relate to the Kon-Notch-Pros glial regenerative gene network, we tested whether loss or gain of function for *Notch* or *pros* might affect *ia-2* expression. With qRT-PCR on dissected larval CNSs, we found that *Notch^ts^* mutants had an almost two-fold increase in *ia-2* expression, whereas *Notch^ICD^* over-expression in glia (*repoGAL4>Notch^ICD^*) caused no significant effect (Figure 1C). Like Kon, Notch also functions in glia (Griffiths and Hidalgo 2004; Kato et al. 2011; Losada-Perez et al. 2016), thus the genetic inference is that *ia-2* expression is prevented by Notch in glia. *Ia-2* mRNA levels also increased (albeit not significantly) in *pros* mutant larvae, but mostly when *pros* was over-expressed in glia (Figure 1D). The loss of function phenotype could be indirect, as in glial cells Pros and Notch depend on each other (Kato et al. 2011), loss of *pros* causes the down-regulation of *Notch*, which would increase *ia-2* expression. Instead, the stronger effect of *pros* gain of function on *ia-2* indicated that Pros could directly regulate *ia-2* expression. Importantly, Pros is a transcription factor found in glia, type I and II neuroblasts, ganglion mother cells and some neurons (Bayraktar et al. 2010). Thus, Pros could regulate *ia-2* expression in any of these cell types. Most importantly, these data meant that *ia-2* participates in the *kon, Notch*, *pros* gene network that drives the regenerative response to CNS injury.

The above data suggested that *ia-2* expression is normally excluded from glia. To test what cells express *ia-2*, we knocked-down *ia-2* with RNAi in either neurons or glia and measured *ia-2* mRNA levels with qRT-PCR in dissected larval CNSs. *ia-2-RNAi* knock-down in glia (*repoGAL4>UASia-2RNAi^TRIPHMS00536^*) had no effect, however knock-down in neurons (*elavGAL4>UASia-2RNAi^TRIPHMS00536^*) down-regulated *ia-2* transcripts to about 20% of wild-type levels (Figure 1E). A second *UAS-ia-2 RNAi* line (line *UAS-ia-2RNAi^KK108555-VIE-260B^*) had a milder effect, but still reduced *ia-2* expression by 25% (Figure 1E). These data meant that *ia-2* is expressed in neurons. To visualize *ia-2* expression *in vivo*, we used a transgenic protein fusion of Ia-2 to yellow fluorescent protein (YFP), Ia-2YFP^CPTI100013^(Lowe et al. 2014; Lye et al. 2014), from now on called Ia-2YFP. Ia-2YFP+ cells did not have the glial marker anti-Repo, nor anti-Deadpan (Dpn), which is the general neuroblast marker and also labels transit amplifying ganglion mother cells in type II neuroblast lineages (Boone and Doe 2008), but all Ia-2YFP+ cells were Elav+ (Figure 1F,G,H). This demonstrated that *ia-2* is expressed exclusively in neurons.

Altogether, these data showed that Ia-2 and Kon function within the regenerative gene network, and are restricted to neurons and glia, respectively (Figure 1I).

### Alterations in Ia-2 levels induce ectopic neural stem cells

Next, we carried out a functional analysis of *ia-2* in the CNS. As *kon* knock-down increased *ia-2* mRNA levels, we sought to verify this using Ia-2YFP. Ia-2YFP+ appeared undistinguishable from wild-type when *kon* was knocked-down in glia (*kon^c452^/ia-2YFP; repoGAL4>kon-RNAi*), but as Ia-2YFP is normally in all neurons, a potential effect could have been missed. Thus, we focused on the midline, where a limited number of dorsal Ia-2YFP+ neurons can be counted. Indeed, *kon* loss of function in glia increased the number of Ia-2YFP+ cells along the midline (Figure 2A,B). The ectopic ia-2YFP cells had the neuronal marker Elav and did not have the glial marker Repo (Figure 2C), meaning they were neurons. Midline cells were unaffected by *kon* over-expression in either neurons or glia (Figure 2A,B, *elavGAL4>kon* and *repoGAL4>kon*). Thus, in the absence of *kon*, ectopic Ia-2YFP+ neurons were found at the midline. Loss of *kon* function prevents glial differentiation (Losada-Perez et al. 2016), and could result in more Ia-2YFP+ neurons also in other locations, but we were unable to verify this. The increase in neurons could explain why *ia-2* mRNA levels increased with *kon* loss of function (see Figure 1A), although the mRNA levels for *ptp99A, -69F* and *10D* (Figure 1 – Figure Supplement 3) also known to function in neurons were not increased. Either way, these data confirmed that Kon and Ia-2 are mutually exclusive in glia and neurons, respectively.

**FIGURE 2.**
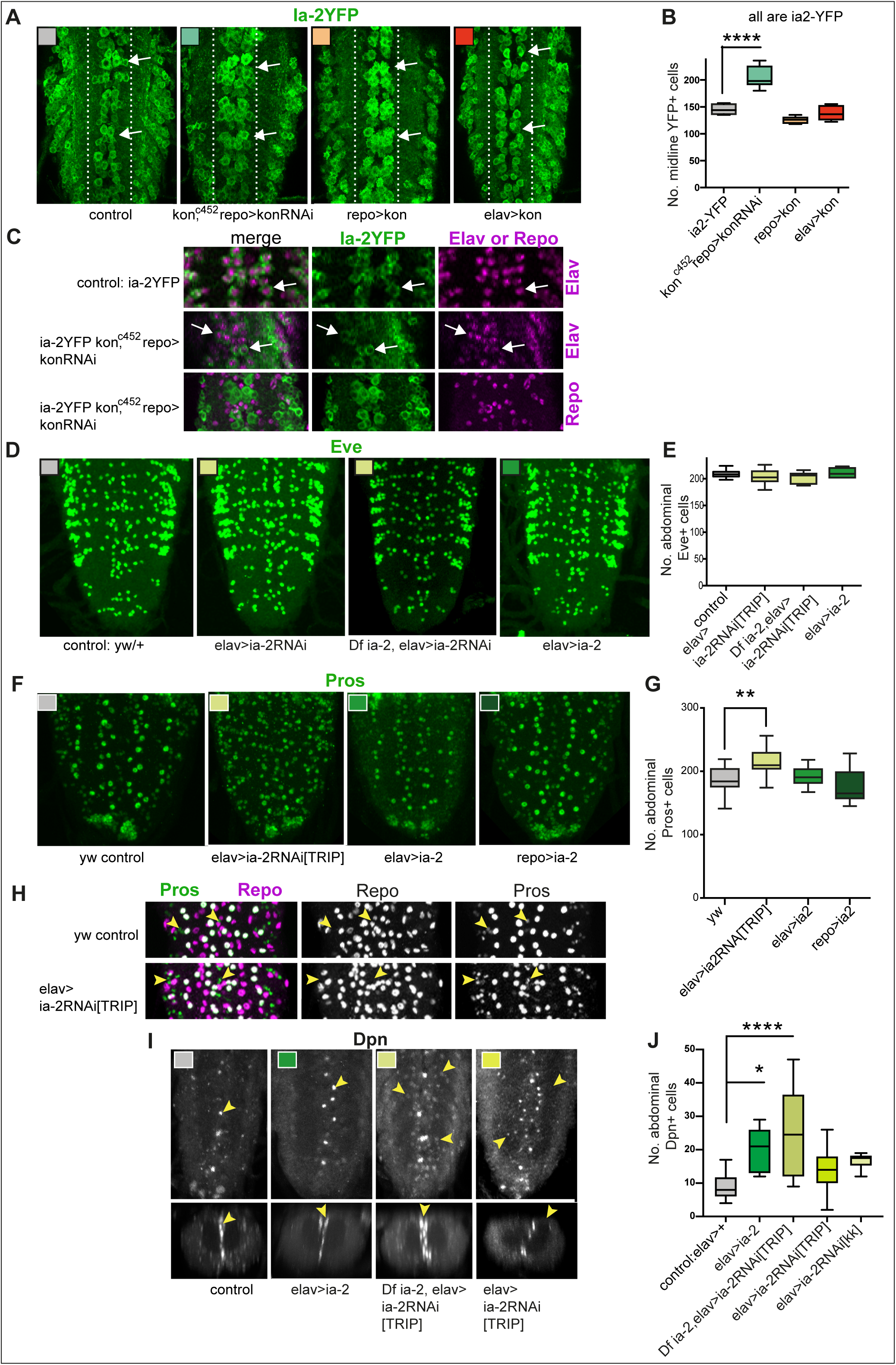
*ia-2* influences neural cell fate non-autonomously. **(A,B)** Loss of *kon* function in glia (*kon^c452^/UASkonRNAi; repoGAL4/+)* increased the number of Ia-2YFP+ cells along the midline. One Way ANOVA p<0.0001, post-hoc Tukey’s test. N=5-8 VNCs. **(C)** The ectopic Ia-2YFP+ cells in *kon* loss of function were Elav+ and not Repo+. N=5-7 VNCs. **(D, E)** Neither loss nor gain *of ia-2* function affected the number of Eve+ neurons. One Way ANOVA p=0. 2374. N=7-12 VNCs. **(F,G)** Loss of *ia-2* function *(elavGAL4>UASia-2RNAi[TRIPHSM00536])* increased Pros+ cell number, and the excess cells were small. Kruskal-Wallis ANOVA p=0.0003, post-hoc Dunnett’s test. N=8-10 VNCs. **(H)** The small Pros+ cells in *ia-2* knock-down did not have the glial marker Repo. **(I,J)** Dpn+ cells visualised at 120h AEL, after developmental neuroblasts have disappeared. Both loss and gain of *ia-2* function increased Dpn+ cell number. All images are horizontal views, except for **(I** bottom row) which are transverse views. One Way ANOVA p=0.0002, post-hoc Dunnett. N=7-15. Asterisks indicate multiple comparison post-hoc tests to a fixed control: *p<0.05, **p<0.01, ***p<0.001, ****p<0.0001. For further statistical analysis details see Table Supplement 1.

To ask what function Ia-2 might have in neurons, we altered *ia-2* expression and visualized the effect using standard neuronal markers. *ia-2* knock-down in neurons (*elavGAL4>ia-2RNAi^TRIPHMS00536^*) had no detectable effects on FasII or BP102 (Figure 2 — Figure Supplement 1A,B), and it did not change Eve+ neuron number either (Figure 2D,E). As Pros activates *ia-2* expression (Figure 1D), we asked whether *ia-2* might affect Pros. Over-expression of *ia-2* in either neurons or glia had no effect on Pros+ cells (Figure 2F,G). By contrast, *ia-2* knock-down in neurons (*elavGAL4>ia-2RNAi ^TRIPHMS00536^*) increased Pros+ cell number, these cells looked small (Figure 2F,G) and lacked the glial marker Repo (Figure 2H). Pros is normally found in ganglion mother cells and neurons, which are generally smaller than glia, suggesting that the ectopic Pros+ cells might be amongst these cell types.

To test whether ectopic Pros+ cells originated from neural stem cells, we asked whether altering *ia-2* function might affect the expression of the general neuroblast marker *dpn*. Both *ia-2* gain of function (*elav>ia-2*) and loss of function (*Df(2L)ED7733/+; elav>ia-2RNAi ^TRIPHMS00536^*, and *elavGAL4* knock-down using two different RNAi lines, *UAS-ia-2RNAi ^TRIPHMS00536^ and UAS-ia-2RNA ^KK108555-VIE-260B^),* in neurons increased the number of abdominal VNC Dpn+ cells (Figure 2I,J). The increase in Dpn+ cell number also correlated with tumorous overgrowths in the VNC (Figure 2 — Figure Supplement 1C), characteristic of genotypes causing neuroblast over-proliferation. The ectopic Dpn+ cells were distributed along the midline, and around the neuropile, in positions normally occupied by glial cells (Figure 2I). The ectopic Dpn+ cells were distinct from normal larval neural stem cells, which are ventro-lateral and further away from the neuropile. Furthermore, they were visualised 120h after egg laying (AEL), after the disappearance of developmental abdominal neural stem cells. Thus, alterations in the levels of neuronal Ia-2 induced neural stem cell marker expression ectopically.

These data showed that interference with normal neuronal Ia-2 levels up-regulated ganglion mother cell and neural stem cell markers. This effect was non-autonomous, as neurons themselves seemed unaffected. As Ia-2 and Kon are functionally related but confined to either neurons or glia, respectively, this suggested that communication between neurons and glia was involved in inducing an ectopic neural stem cell state.

### Injury induces *ia-2* expression and a regenerative neurogenic response

Above data had shown that altering *ia-2* levels up-regulated the neural stem cell marker Dpn (Figure 2I,J). Since CNS injury induced the up-regulation of *kon* expression (Losada-Perez et al. 2016), we asked whether injury might affect *ia-2* expression and, consequently, induce a neurogenic response. To this end, crush injury was carried out at 74-76h after egg laying (AEL) in early third-instar larval VNCs labelled with the endoplasmic reticulum GFP marker G9 (Figure 3A,D), using a previously established protocol (Losada-Perez et al. 2016). qRT-PCR in injured VNCs revealed approximately a 2-fold increase in *ia-2* mRNA levels at 5-7h post-injury, which recovered homeostatically by 24h post-injury (Figure 3D). This paralleled the effect of injury on *kon* expression (Losada-Perez et al. 2016). Thus, CNS injury caused an increase in *ia-2* expression.

**FIGURE 3.**
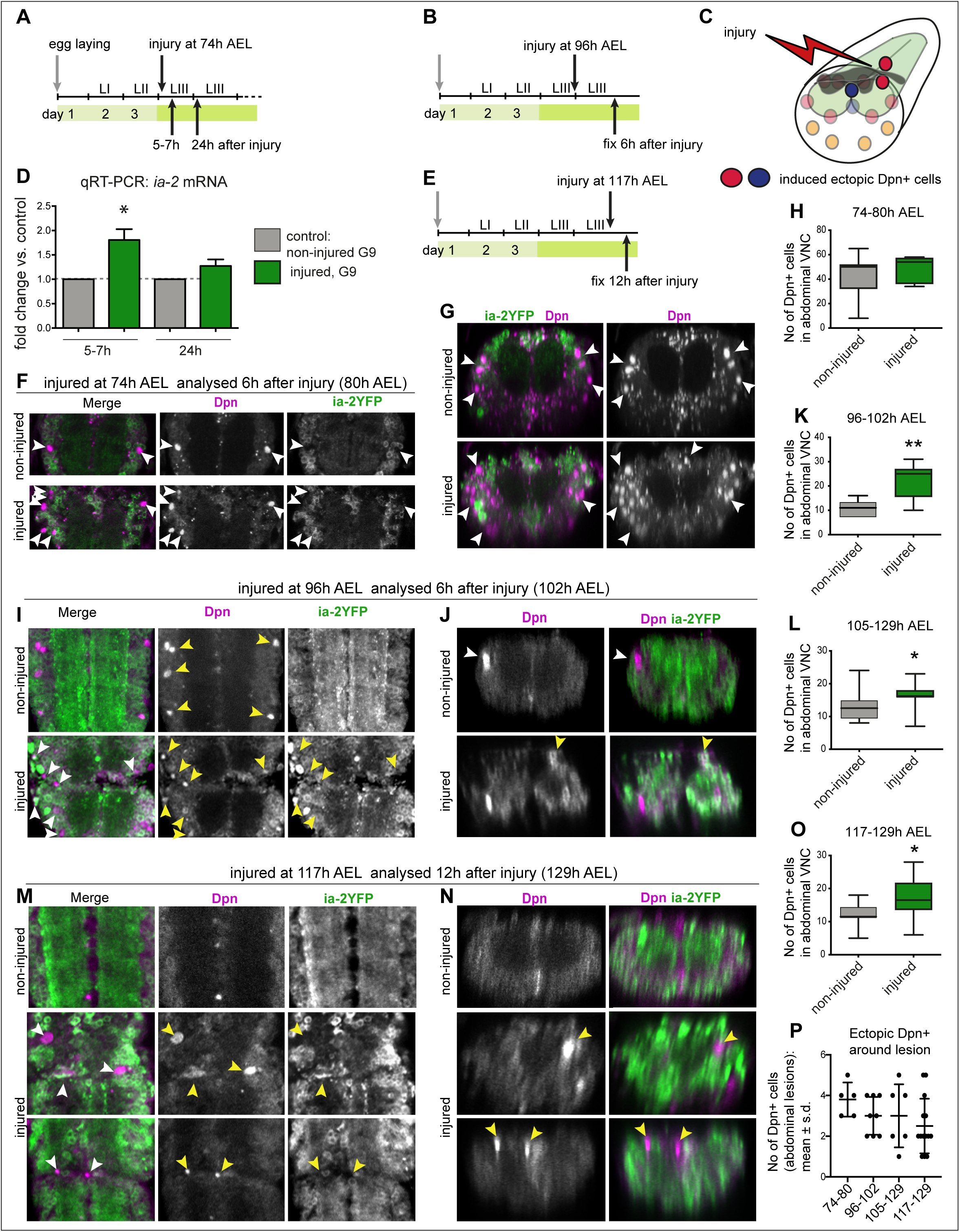
Injury induced *ia-2* expression and ectopic Dpn+ cells. **(A,B,E)** Time course of crush-injury experiments, indicating the age of the larvae (After Egg Laying, AEL) when crush was applied (top arrows), followed by various recovery periods, and when they were fixed or processed (bottom arrows). **(C)** Drawing showing that crush injury induced ectopic Dpn+ cells. **(D, F,G,H)** Crush injury in the larval abdominal VNC at 74-76h after egg laying (AEL) caused: **(D)** an increase in the levels of *ia-2* mRNA at 5-7h post-injury, which recovered homeostatically by 24h, detected by qRT-PCR. N=3 replicates. **(F,G,H)** Formation of ectopic Dpn+ neural stem cells (white arrowheads) by 5-7 hours post-injury, 74-80h AEL. Quantification in **(H)** shows number of Dpn+ in VNCs with ectopic Dpn+ cells (Penetrance 50% N=10 VNCs), as in some VNC cell loss caused by lesion was very severe. Dpn+ cells were Ia-2YFP^—^. **(I,J,K)** Crush injury in the larval abdominal VNC at 96h AEL caused ectopic Dpn+ cells by 6 hours post-injury (yellow arrowheads, penetrance 44% N=9 VNCs). Most Dpn+ cells were Ia-2YFP^—^, but some were Ia-2YFP^+^. At this stage, some developmental neuroblasts could still remain (white arrows), but dorsal ectopic Dpn+ were unequivocal (yellow arrowheads, **J**). (**K)** Student t-test p=0.0063. **(L)** Injury at 105h AEL visualised at 129h AEL, when no developmental neuroblasts remain, induced a significant increase in Dpn+ cells. Mann-Whitney U-test p=0.0375. **(M,N,O)** Crush injury in the larval abdominal VNC at 117h AEL caused ectopic Dpn+ cells by 12 hours post-injury (yellow arrowheads, 129h AEL, Penetrance 67% N=21, VNCs. Student t-test p=0.0302). Dpn+ cells were found in ectopic dorsal positions (yellow arrowheads, **N**). This stage is devoid of developmental neural stem cells. N=9/32 VNCs. Student t-test. **(P)** Temporal profile of number of ectopic Dpn+ cells surrounding the lesions, in injured samples with ectopic Dpn+ cells, number in X axis indicate time-points of injury and fixation. **(F,I,M)** Horizontal views, **(G,J,N)** transverse views. **(H,K,L,O)** Graphs show quantifications in box-plots; **(P)** shows dot plots, with mean and error bars (±s.d.) indicated. *p<0.05, **p<0.01. For full genotypes and further statistical analysis details see Table Supplement 1.

Since increased *ia-2* levels induced ectopic Dpn+ cells (Figure 2I,J), and *ia-2* was up-regulated in injury, we asked whether injury induced neural stem cells. We focused in the abdominal VNC only, which has 3 neuroblasts per hemi-segment in ventro-lateral positions, in early third instar larvae. Crush injury in the abdominal VNC at 74-76h AEL resulted in ectopic Dpn+ cells by 5-7h later (Figure 3A,F,G,H). These were more numerous than the normal developmental abdominal larval neuroblasts, and included cells located in dorsal positions, which are not normally occupied by them (Figure 3F,G; see (Sousa-Nunes et al. 2011; Froldi et al. 2015). The numerous Dpn+ cells could correspond to injury-induced divisions of neuroblasts normally found during larval development. To test whether injury might induce ectopic neural stem cells distinct from developmental neuroblasts, we next carried out crush injury at three later time points: (1) at 96h AEL and analysed the VNCs 6h post-injury (PI, 102h AEL), when in control VNCs, abdominal hemi-segments have 0 or 1 Dpn+ cells remaning (Figure 3B,I,J,K); (2) At 105h and analysed 24h PI (129h AEL), when in controls there are no ventro-lateral neuroblasts, only Dpn+ cells along the midline (Figure 3L); and (3) at 117h AEL and analysed the VNCs 12h PI (129h AEL), taking advantage of the delayed pupariation of injured larvae (Figure 3M,N,O). At 129h AEL there were no remaining abdominal ventro-lateral neural stem cells in intact controls, only some Dpn+ cells along the midline (Figure 3N). Injury induced at these three later time points, also caused ectopic Dpn+ cells compared to controls (Figure 3K,L,O), and most, if not all, ectopic Dpn+ cells lacked Ia-2YFP (Figure 3F,G,I,J,M,N). Importantly, most ectopic Dpn+ cells surrounded the neuropile, and some were dorsal, in positions never occupied by developmental neural stem cells (Figure 3J,N). In the samples with ectopic Dpn+ cells, these were found surrounding the lesions (Figure 3I-N,P). These data showed that injury induces ectopic neural stem cells, and these are distinct from developmental neuroblasts. Since *ia-2* levels increased upon injury, and *ia-2* gain of function induced neural stem cells, this suggested that *ia-2* was responsible for the increase in Dpn+ cells caused by injury.

### Dilp-6 depends on neuronal Ia-2 and glial Kon

The above data raised the question of how Ia-2 might induce ectopic neural stem cells. Ia-2 is highly evolutionarily conserved and it functions in dense core vesicles to release insulin and neurotransmitters (Harashima et al. 2005; Kim et al. 2008; Nishimura et al. 2010; Cai et al. 2011). There are eight *Drosophila* insulin-like-peptides (Dilps) and Ia-2 affects only Dilp-6 (Kim et al. 2008). *dilp-6* is expressed in cortex and blood brain barrier CNS glia, and activates neural stem cell proliferation following a period of quiescence in normal larval development(Sousa-Nunes et al. 2011) (Chell and Brand 2010; Sousa-Nunes et al. 2011). Thus, we asked whether the increase in Dpn+ cells in *ia-2* loss and gain function observed above involved *dilp-6*.

We first visualized *dilp-6* expressing cells in wandering larvae using *dilp6-GAL4* (Chell and Brand 2010; Sousa-Nunes et al. 2011) to drive expression of the nuclear reporter Histone-YFP (His-YFP). Most *dilp-6>his-YFP+* cells were also Repo+, but they did not surround the neuropile and lacked the neuropile glial marker Pros (Figure 4A,B). Therefore, most *dilp-6* expressing cells in the abdominal larval VNC were cortex and surface glia, as previously reported (Chell and Brand 2010; Sousa-Nunes et al. 2011). Some *dilp6>his-YFP+* cells were Repo— Elav+, and thus were neurons (Figure 4A,B). Therefore, *dilp-6* is expressed in some neurons per VNC segment, and mostly in non-neuropile glia.

**FIGURE 4.**
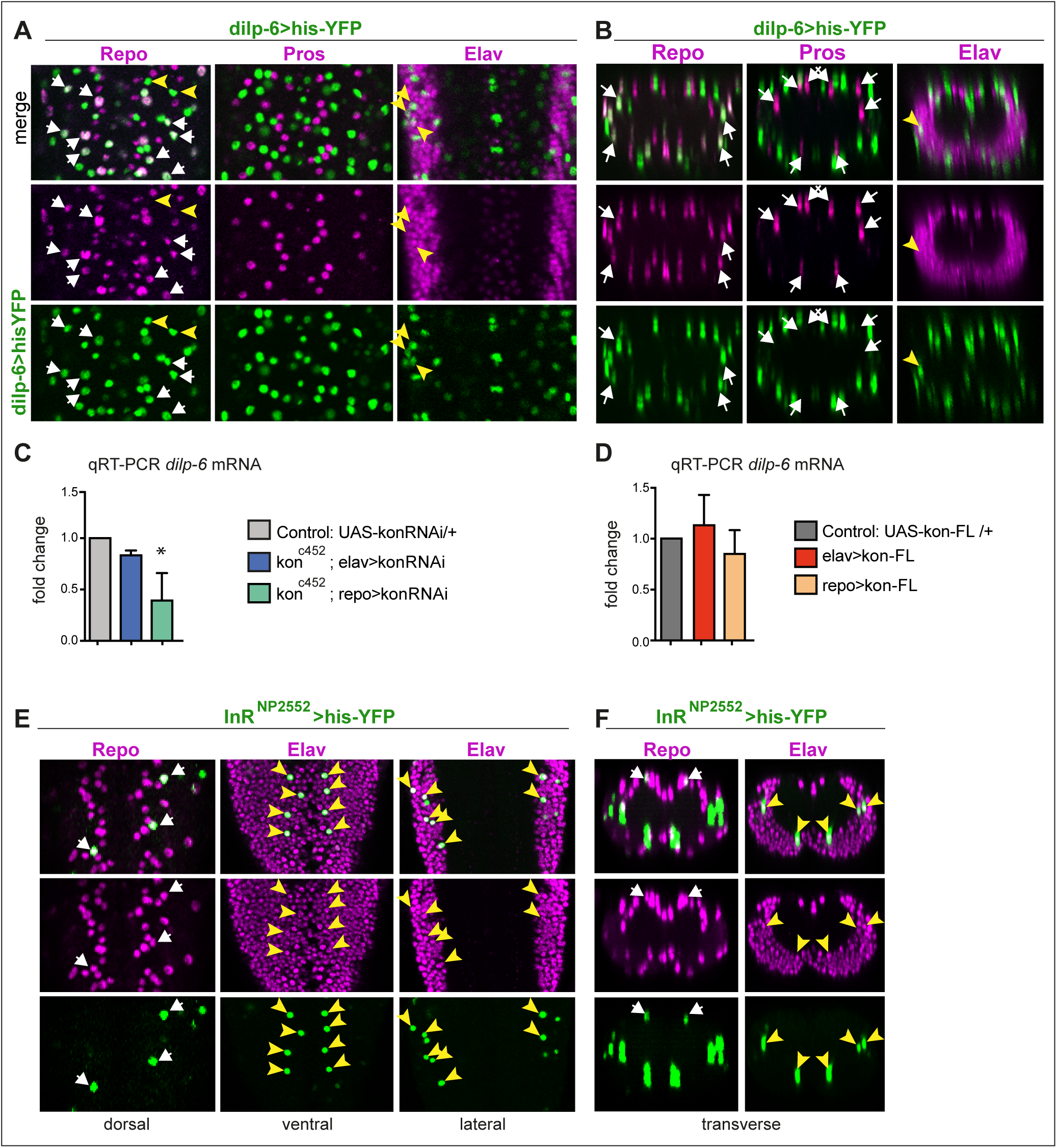
*dilp-6* is expressed in neurons and cortex glia and received by neuropile glia. **(A,B)** *Dilp-6GAL4>UAShisYFP* cells are mostly Repo+ Pros^—^glia that do not surround the neuropile (white arrows), and from position appear to be cortex and surface glia. No YFP+ cells have Pros. Some cells are Repo^—^ Pros^—^ Elav+ (yellow arrowheads) meaning they are neurons. **(C,D)** qRT-PCRs showing that: **(C)** *kon* knock-down in glia *(kon^c452^/UASkonRNAi; repoGAL4/+)* downregulates *dilp-6* mRNA levels; **(D)** over-expression on *kon* does not cause a significant effect. N=3 replicates for both. **(C,D)** One Way ANOVA, only differences in **(C)** for *dilp-6* mRNA significant p=0.0362, *p<0.05. **(E,F)** *inR* expression visualized with reporter *InR^NP2552^GAL4> UAShistoneYFP* is expressed stochastically in some dorsal Repo+ neurople glia (white arrows), and other glia, and in some Elav+ neurons (yellow arrowheads). **(A,E)** Horizontal views of the abdominal VNC; **(B,F)** transverse views. For full genotypes and further statistical analysis details see Table Supplement 1.

Kon function is required for glial cell fate (Losada-Perez et al. 2016), so we used qRT-PCR to ask whether altering Kon levels might affect *dilp-6* expression. Over-expression of full-length *kon* midly increased *dilp-6* mRNA levels (albeit not significantly), but *kon* knock-down in glia significantly reduced *dilp-6* mRNA levels (Figure 4C,D). This effect could be indirect, as glial proliferation and differentiation are impaired with *kon* loss of function (Losada-Perez et al. 2016), or perhaps Kon regulates *dilp-6* expression. Either way, *dilp-6* expression depends on *kon* in glia.

Dilp-6 is a ligand for the *insulin receptor* (*InR*), which is expressed and/or functional at least in CNS neurons and neuroblasts (Fernandez et al. 1995; Song et al. 2003; Sousa-Nunes et al. 2011; Fernandes et al. 2017). To revisit what cells might receive Dilp-6, we visualized *InR* expression, using available GAL4 lines to drive *his-YFP,* and tested co-localisation with glial and neuronal markers. At 72h AEL, *InR^NP2552^>his-YFP+* cells comprised some Elav+ neurons and some Repo+ glia, including dorsal neuropile glia and surface glia (Figure 4E,F). The distribution was stochastic, most likely due to the insertion of GAL4 into an intron. According to these data limited by the currently available tools, *InR* is expresed in both neurons and glia.

Altogether, these data indicated that in the third instar larva Dilp-6 is produced and secreted by Ia-2 in some neurons, it is mostly produced in non-neuropile glia, and it is received by InR in neurons and glia, including neuropile glia.

### A positive neuron-glia communication loop boosts Dilp-6 production from glia

The above data strongly suggested that a neuron-glia communication loop might serve to amplify Dilp-6. A limiting step could be Kon, as glial *dilp-6* expression depends on *kon*. Kon is required for glial gene expression (Losada-Perez et al. 2016), but whether this depends on the nuclear translocation of its intracellular domain, Kon^ICD^, is unknown. In *Drosophila*, Kon had been reported to lack a nuclear localization signal (Schnorrer et al. 2007). In mammals, NG2^ICD^ positively regulates the expression of multiple genes, including downstream targets of mTOR (Sakry et al. 2015; Nayak et al. 2018), but whether this requires nuclear NG2^ICD^ is also unknown. Altogether, whether NG2 or Kon regulate glial gene expression through nuclear events remained unsolved. Thus, to ask whether Kon^ICD^ might function in the nucleus, we generated an HA-tagged form of Kon^ICD^ (Kon^ICD-HA^). Glial over-expression of *kon^ICD-HA^* (*repoGAL4>UAS-Kon^ICD-HA^*) revealed distribution of anti-HA in glial cytoplasms and co-localisation with the glial nuclear transcription factor Repo in glia cells, both in embryos and larvae (Figure 5A and Figure 5 – Figure Supplement 1)). Thus, Kon^ICD^ is distributed in the cytoplasm and nucleus, from where it could regulate gene expression.

**FIGURE 5.**
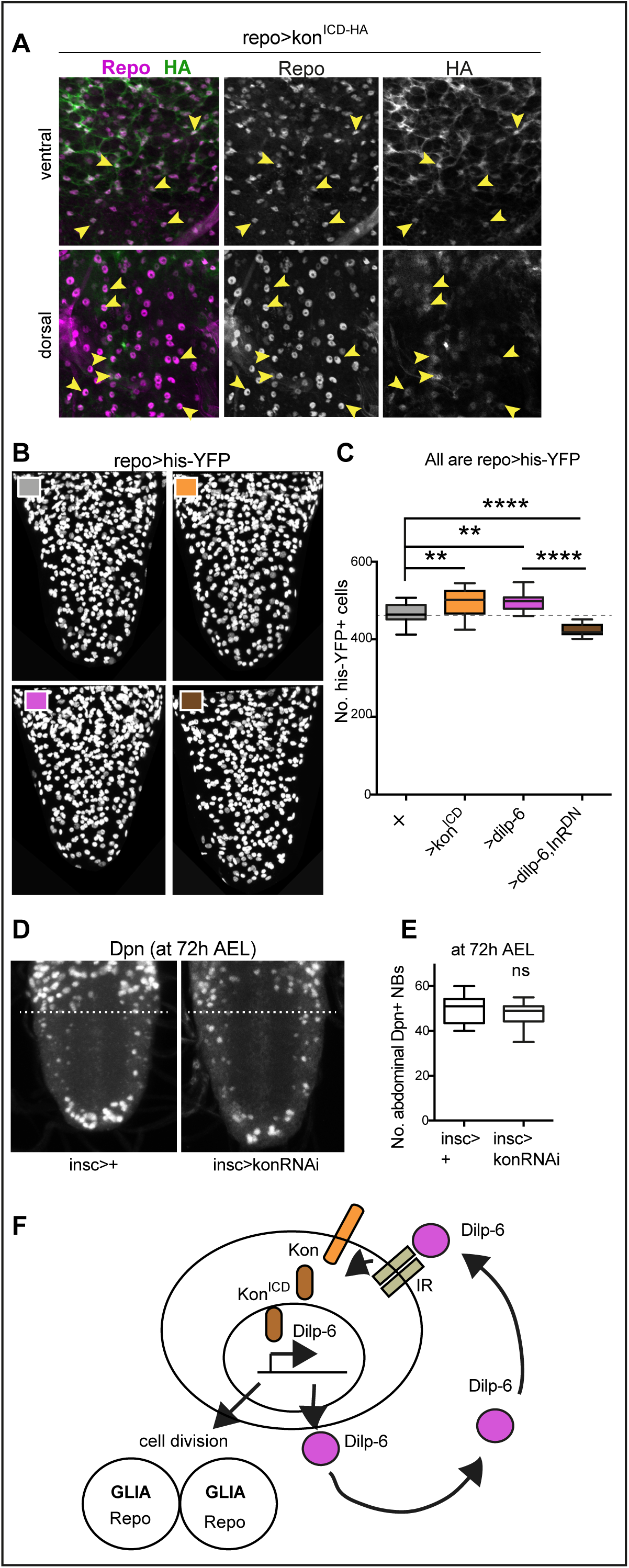
Ia-2, Kon and Dilp-6 are linked though a neuron-glia communication loop. (A) Over-expressed HA-tagged Kon^ICD^ in glia (*repoGAL4>UASkon^ICd^::HA*) visualized with anti-HA antibodies in third instar wandering larvae, localizes to both glial cytoplasms and nuclei (arrows). **(B,C)** Over-expression of either the intracellular domain of either *kon (kon^ICD^)* or *dilp-6* increased glial cell number, visualized with *repoGAL4>UAShistone-*YFP, and quantified automatically with DeadEasy in **(C).** Over-expression of a dominant negative form of the insulin receptor rescues the increase in cell number caused by Dilp-6 *(repo>hisYFP, dilp-6, InRDN),* meaning that autocrine InR signaling regulates glial proliferation. One Way ANOVA p<0.0001, post-hoc Tukey’s test multiple comparisons between all samples. N=15-28 VNCs. **(D,E)** Third star larvae at 72h AEL to visualise abdominal developmental neuroblasts: *kon-RNAi* knock-down in neural stem cells with *inscGAL4* does not affect Dpn+ cell number. Unpaired Student t-test, p=0.3111. N=10 VNCs **(F)** Illustration summarising that a positive feedback autocrine loop involving Dilp-6, InR and Kon promotes both glial proliferation and Dilp-6 production. All images are horizontal views. Asterisks refer to multiple comparison post-hoc tests, all samples vs. all: **p<0.01, ****p<0.0001. All graphs show box-plots. For full genotypes and further statistical analysis details see Table Supplement 1.

Next, we asked whether Kon^ICD^ is functional. Since NG2 and Kon are responsible for glial proliferation both in mammals and *Drosophila* (Kucharova and Stallcup 2010; Losada-Perez et al. 2016), we used glial cell number as a read-out of Kon^ICD^ function. First, we tested whether cleaved Kon^ICD^ could induce glial proliferation, like full-length Kon does (Losada-Perez et al. 2016). We over-expressed *kon^ICD^* in glia and automatically counted glial cells labeled with the nuclear marker his-YFP, using DeadEasy software (Forero et al. 2012). Over-expression of *kon^ICD^* in glia increased glial cell number (*UAShisYFP; repoGAL4>UASkon^ICD^*, Figure 5B,C), meaning that Kon^ICD^ can induce glial proliferation. As full-length Kon also promotes glial proliferation (Losada-Perez et al. 2016), these data meant that full-length Kon is normally cleaved, releasing Kon^ICD^ to promote glial proliferation, although we were unable to verify whether it regulates gene expression.

In principle, Dilp-6 amplification could occur if it were first secreted from neurons by Ia-2, to activate InR in glia, and InR signalling in turn drove the Kon-dependent up-regulation of *dilp-6* expression in glia (Figure 5F). To test whether Dilp-6 activates InR in glia, which activates Kon, we asked: (1) whether over-expression of *dilp-6* could mimic the increase in glial cell number caused by Kon^ICD^, and (2) whether this could be rescued by over-expression of a dominant negative form of the *insulin receptor (InR^DN^)* in glia. We found that over-expression of *dilp-6* in glial cells increased glial cell number comparably to Kon^ICD^ (Figure 5B,C), and this was rescued with concomitant over-expression of *InR^DN^* in glia (Figure 5B,C). These data meant that Dilp-6 activates InR signaling in glia, and induces glial proliferation.

Dilp-6 and InR signaling reactivate quiescent developmental neural stem cells (Chell and Brand 2010; Sousa-Nunes et al. 2011), but Kon functions in glia (Losada-Perez et al. 2016). To further verify whether Kon function is restricted to glia, we asked whether Kon might also be required in neural stem cells during development at 72h AEL, when there normally are neural stem cells in both thorax and abdomen of larvae. RNAi *kon* knock-down in neural stem cells with *inscutable-GAL4 (ins-GAL4>UAS-konRNAi)* did not affect the number or distribution of abdominal developmental Dpn+ cells at 72h AEL (Figure 5D,E), meaning that Kon is not required for neural stem cell development. Since glial proliferation depends on Kon (Losada-Perez et al. 2016), the fact that *dilp-6* alone could reproduce the increase in cell number caused by *kon^ICD^,* and this depended on InR in glia, strongly suggested that InR signalling can activate Kon cleavage downstream in glia.

To conclude, altogether these data suggested that Ia-2 triggers the release of Dilp-6 from neurons, which then is received by glial cells, where InR signaling activates Kon, which in turn induces glial proliferation, and further expression of *dilp-6*. Thus, a non-autonomous relay from neuronal Ia-2 to glial Kon promotes glial proliferation and induces a positive feedback loop that amplifies Dilp-6 production from glia (Figure 5F).

### Ia-2 and Dilp-6 can induce neural stem cells from glia

So far, our data had shown that: alterations in Ia-2 levels caused either by genetic manipulation or injury induced ectopic neural stem cells; Ia-2 is required for the neuronal secretion of Dilp-6, which is received and amplified in cortex glia under the control of Kon; and secreted Dilp-6 is received by InR also in neuropile glia. As Dilp-6 activates quiescent developmental neural stem cells (Chell and Brand 2010; Sousa-Nunes et al. 2011), this raised the question of whether the Ia-2 -Kon-Dilp-6 loop not only produced more glia, but could also induce a neurogenic response from glia.

To ask whether Kon, Ia-2 or Dilp6 could be responsible for inducing ectopic neural stem cells from glia, we over-expressed them in glia (with *repoGAL4*), and analysed Dpn at 120h AEL, after the disappearance of developmental abdominal neural stem cells. Over-expression of *kon* did not induce ectopic Dpn+ cells (Figure 6A-D). By contrast, over-expression of *ia-2* induced ectopic Dpn+ cells prominently along the midline but also in lateral locations surrounding the neuropile, ordinarily occupied by glia (Figure 6A-D). Over-expression of *dilp-6* had a stronger effect, and there were many ectopic Dpn+ cells surrounding the neuropile (Figure 6A-D). Dpn levels in ectopic cells were generally lower than in normal neural stem cells. These data showed that both Ia-2 and Dilp-6 can induce *dpn* expression, potentially in glia. However, Kon alone cannot, meaning that insulin signaling is required to induce neural stem cells. Since Ia-2 drives Dilp-6 production and secretion, this suggested that ultimately Dilp-6 induced ectopic neural stem cells (Figure 6E).

**FIGURE 6.**
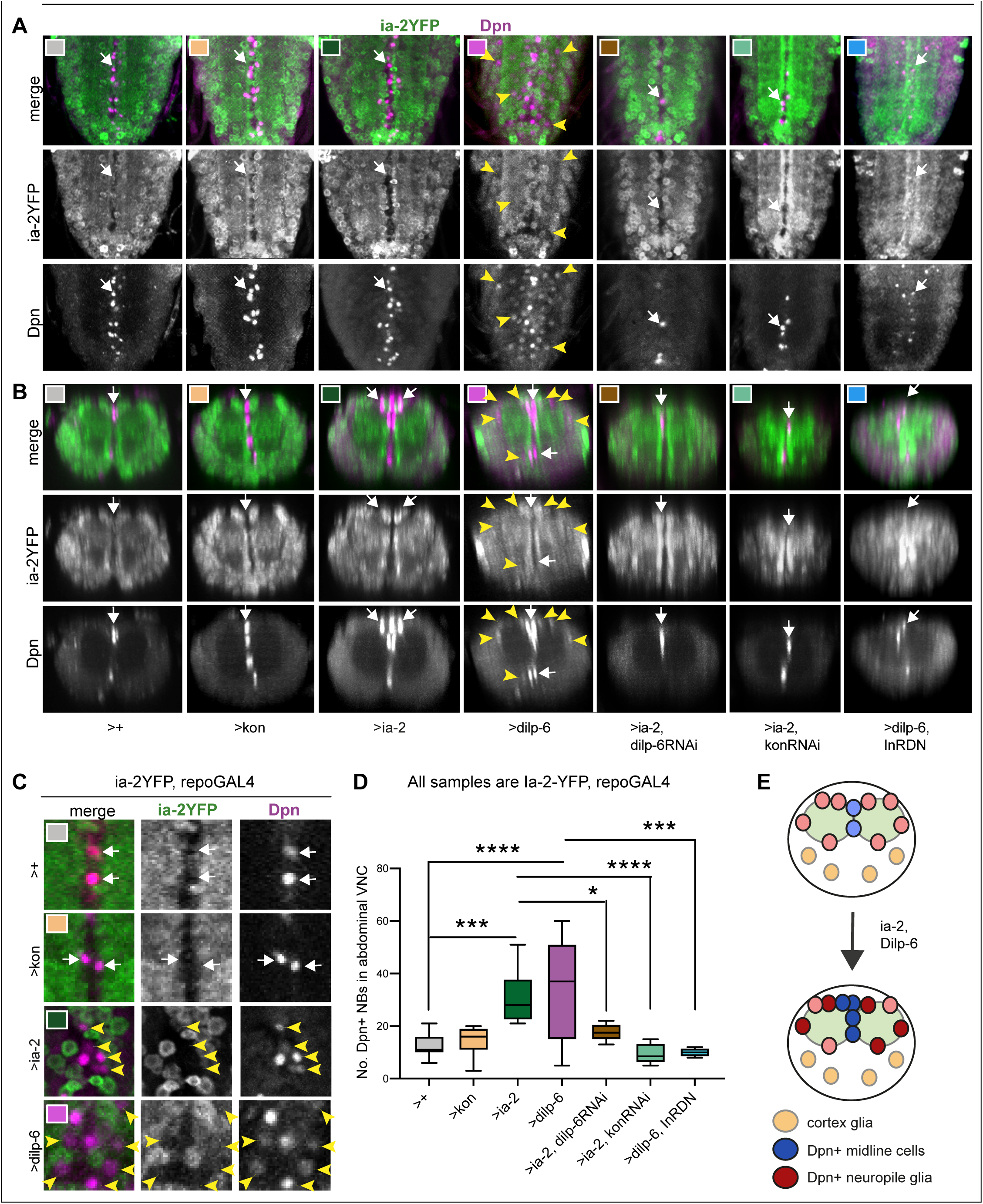
Ia-2 and Dilp-6 induce ectopic neural stem cells from InR signaling in glia. All samples were analysed at 120h AEL, after disappearance of abdominal developmental neuroblasts. **(A-C)** Over-expression of *ia-2* and *dilp-6*, but not *kon-full-length*, increased Dpn+ cell number in the abdominal VNC. Both Ia-2 and Dilp-6 induced Dpn+ at the midline and in lateral positions: *ia-2* most prominently, but not exclusively, along the midline (white arrowhead), and *dilp-6* also prominently, but not exclusively, in lateral positions around the neuropile (yellow arrowheads). **(C)** Ectopic Dpn+ cells did not express Ia-2YFP (arrowheads). **(D)** Quantification of all abdominal VNC Dpn+ cells, and genetic epistasis analysis showing that: the increase in Dpn+ cell number caused by *ia-2* over-expression was rescued by *dilp-6 RNAi* and *kon-RNAi* knock-down in glia, meaning that *ia-2* requires Dilp-6 and glial Kon to induce Dpn; and preventing insulin signaling with InR^DN^ in glia rescued the increase in Dpn+ cell number caused by *dilp-6* over-expression, meaning that Dilp-6 induced Dpn via InR signalling in glia. One Way ANOVA p<0.0001, post-hoc Tukey’s test multiple comparisons all samples vs. all. N=7-13 VNCs. **(E)** Illustration showing that Ia-2 and Dilp-6 can induce Dpn via InR signalling in glial cells. **(A)** Horizontal views; **(B)** transverse views; **(C)** higher magnification. Graphs shows quantifications in box-plots. Asterisks refer to multiple comparison post-hoc tests: *p<0.05, ***p<0.0001, ****p<0.0001. For full genotypes and further statistical analysis details see Table Supplement 1.

To further test whether Ia-2 up-regulated *dpn* ectopically via Dilp-6, we carried out epistasis analysis. Over-expression of *ia-2* together with *dilp-6* knock-down in glia (*repoGAL4>UAS-ia-2, UAS-dilp-6RNAi)*, rescued the number of Dpn+ cells (Figure 6A-D), demonstrating that Ia-2 induces ectopic neural stem cells via Dilp-6. Furthermore, over-expression of *ia-2* together with *kon* RNAi in glia (*repoGAL4>UAS-ia-2, UAS-konRNAi*) also rescued the Dpn+ phenotype (Figure 6A-D), confirming that *dilp-6* expression depends on Kon in glia (see Figure 4C) and that Kon and Dilp-6 engage in a positive feedback loop (see Figure 5). Finally, the ectopic Dpn+ phenotype was also rescued by over-expression of *dilp-6* together with *InR^DN^* in glia (Figure 6A-D, *repoGAL4>UAS-dilp6, UAS-InR^DN^*), meaning that ectopic neural stem cells depend on InR signaling in glia. Together, these data showed that Ia-2 induces ectopic neural stem cells via Dilp-6 and InR signalling in glia, and that ectopic Dpn cells originated from glia (Figure 6E).

To further test whether the ectopic Dpn+ cells originated from glia, we first asked whether ectopic Dpn colocalised with the glial marker Repo, analysing larvae at 120h AEL, after the disappearance of developmental abdominal neural stem cells. Lateral ectopic Dpn+ cells observed with *dilp-6* over-expression were also Repo+ (Figure 7A,B), consistent with originating from glial cells. Dpn levels were lower than in normal neural stem cells. By contrast, the ectopic midline Dpn+ cells were not Repo+. Midline glia do not normally express *repo*, but express *wrapper (wrp).* Over-expression of *dilp-6* resulted in Dpn+ cells along the midline that also had Wrp (Figure 7C,D), showing that the ectopic midline Dpn+ cells were midline glia. These data showed that there are two distinct populations of ectopic Dpn+ cells: latero/dorsal Repo+ cells around the neuropile and midline Wrp+ cells, altogether indicating that Dpn is induced in neuropile glia (class known as ‘astrocytes’) and midline glia.

**FIGURE 7.**
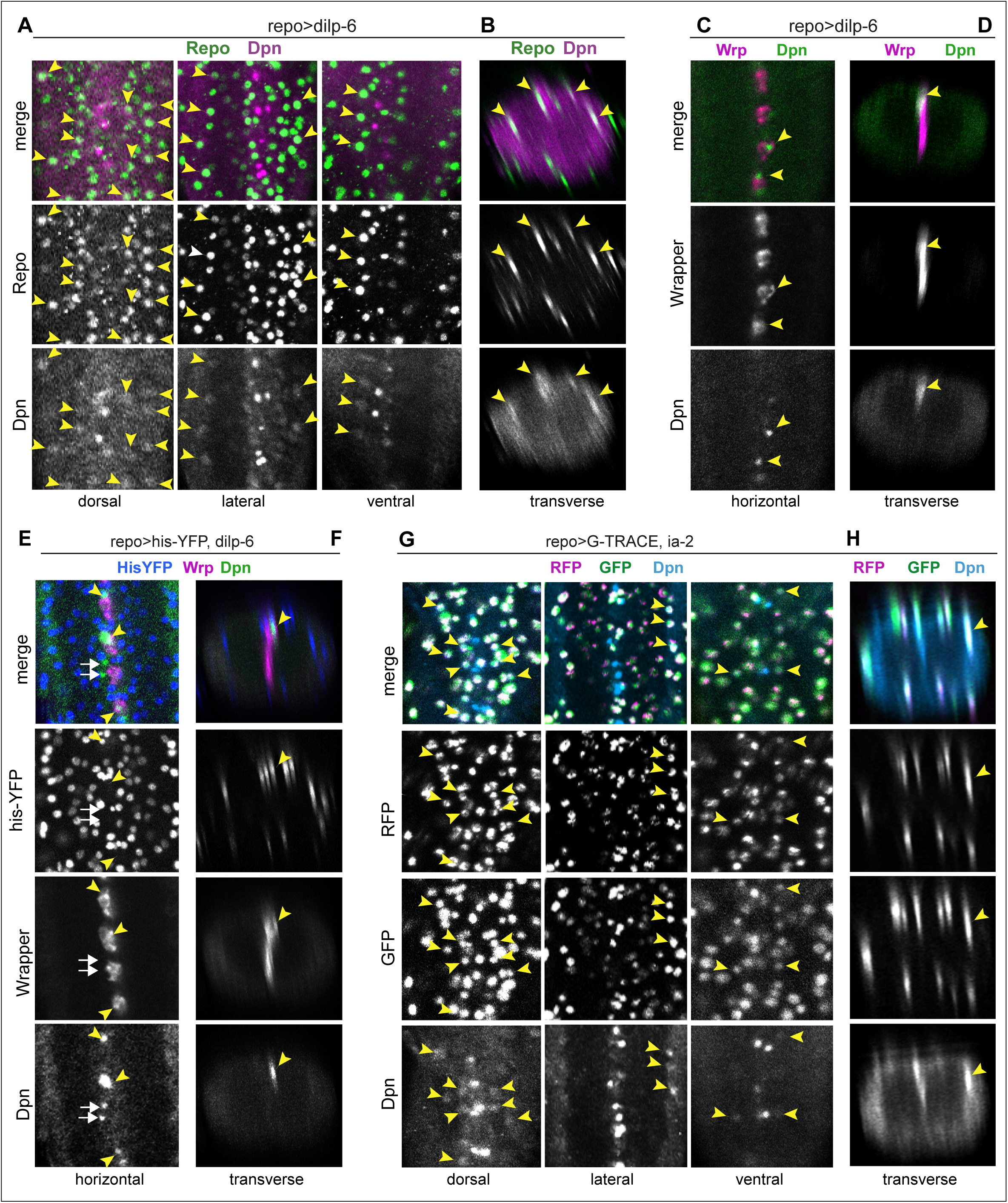
Ia-2 and Dilp-6 induced ectopic neural stem cells originate from glia. All samples were analysed at 120h AEL, after disappearance of abdominal developmental neuroblasts. **(A,B)** Over-expression of *dilp-6* from glia *(repoGAL4>UAS-dilp-6)* induced Dpn expression in Repo+ neuropile glial cells (arrowheads). N= 10 VNCs. **(C,D)** Over-expressed *dilp-6* also induced Dpn in Wrp+ midline glia (arrowheads). N=6 VNCs. **(E,F)** When *dilp-6* was over-expressed, and all glia except midline glia were visualised with nuclear *repoGAL4>Histone-YFP* and midline glia with anti-Wrp, Dpn+ YFP—Wrp— cells were found, which therefore were not glia (white arrows; yellow arrowheads point to Dpn+Wrp+ cells). N=6 VNCs. **(G,H)** G-TRACE expression in glia with *repoGAL4* revealed with GFP cells that were originally glia or originated from a glial cell lineage, even if they switched off the glial *repo* promoter, and with RFP newly generated glial cells. Dpn colocalised in neuropile glia with both GFP and RFP, meaning that Dpn+ cells originated from glia, and at that point in time these cells still retained active the glial *repo* promoter. N= 8 VNCs. **(A,C,E,G)** horizontal and **(B,D,F,H)** transverse views. For full genotypes and sample sizes see Table Supplement 1.

However, not all Dpn+ cells were Repo+ or Wrp+, as some did not express either of these markers (Figure 7E, white arrows; genotype: *repoGAL4>his-YFP, dilp-6*). This could mean that either some ectopic Dpn+ did not originate from glia, or that as glial cells reprogrammed into neural stem cells, they switched off glial gene expression. To test whether ectopic neural stem cells originated from glia, we used the cell-lineage marker G-TRACE. This GAL4-dependent tool results in the permanent labelling of GAL4/UAS-expressing cells and their lineage. Thus, as glial cells become neural stem cells, the glial *repo* promoter would be switched off, but G-TRACE would enable their visualisation as well as that of all their progeny cells. Cells that were originally glia but may no longer be so would be labelled in green (GFP+), and recently specified glial cells would be labelled in red (RFP+). G-TRACE expression in glia with *repoGAL4* together with *dilp-6* caused larval lethality and thus could not be analysed. By contrast, over-expression of both G-TRACE and *ia-2* in glia (*repoGAL4>G-TRACE, UAS-ia-2*) revealed G-TRACE+ Dpn+ cells around the neuropile, at 120h AEL (Figure 7G,H). Most, if not all, of these cells had GFP, but also RFP (Figure 7G,H). These data demonstrate that ectopic Dpn+ originate from glial cells. Since RFP was also present, this suggests that glial cell fate had not been suppressed, and instead that glial cells may have been in the process of reprogramming.

Altogether, these data showed that Ia-2 and Dilp-6 can induce de novo formation of neural stem cells from neuropile and midline glial cells.

### In vivo reprogrammed glial cells can divide and generate neurons

To ask whether ectopic neural stem cells can divide to generate neurons, we used the S-phase marker PCNA-GFP, and we over-expressed *dilp-6* specifically at the third instar larva using GAL4 under the control of a heat-shock promoter. We heat-shocked larvae at 110.5h AEL at 37°C for 30 minutes, then kept them at 25°C for 9 hours, when they were dissected and fixed, to visualize Dpn+ and PCNA-GFP at 120h AEL. In control wandering third instar larvae, a few PCNA-GFP + cells could be observed along the midline, but not in lateral positions (Figure 8A,A’). Over-expressed *dilp-6* resulted in ectopic Dpn+ PCNA-GFP+ cells in lateral positions around the neuropile (Figure 8B,B’,D), and along the midline (Figure 8C,C’,D). Furthermore, Dilp-6 also resulted in PCNA-GFP+Wrp+ cells along the midline (Figure 8E-H), and in an increase in Wrp+ cells (Figure 8G). Finally, over-expression of *ia-2* also increased cell division in midline cells visualised using the mitotic marker anti-phospho-Histone-H3, albeit not significantly (Figure 8I-L). These data demonstrate that Ia-2 and Dilp-6 glial-reprogrammed neural stem cells can divide.

**FIGURE 8.**
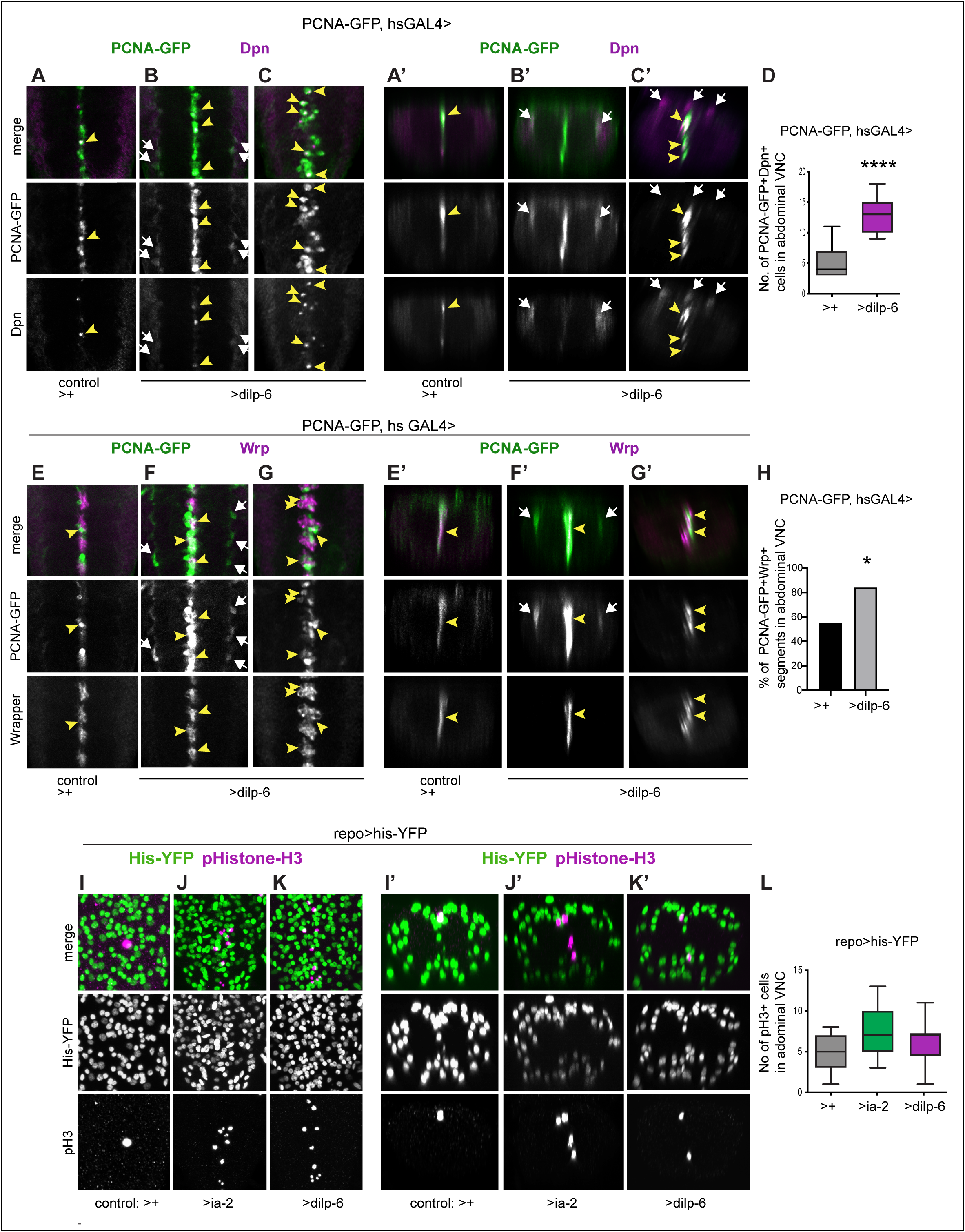
Ia-2 and Dilp-6 induced ectopic neural stem cells can divide. All samples were analysed at 120h AEL, after disappearance of abdominal developmental neuroblasts. **(A-C’, E-G’)** Cell proliferation was visualised with the S-phase marker PCNA-GFP, quantification in **(D,H).** *dilp-6* expression was induced in all cells with *heat-shock-GAL4*, raising the temperature to 37°C for 30 minutes at the end of the third instar larval stage at 110.5h AEL, then larvae were kept at 25°C for 9 hours, visualizing Dpn+ and PCNA-GFP at 120h AEL. **(A-C’)** Over-expression of *dilp-6* resulted in Dpn+ PCNA-GFP+ cells laterally around the neuropile (**B,B’** white arrows) and along the midline (**C,C’** yellow arrowheads), showing that these ectopic Dpn+ cells were in S-phase. Quantification box-plots in **(D)**, Student t-test. There were also some Dpn+ cells that were not dividing (white arrows in C’). **(E-G’)** Over-expression of *dilp-6* resulted in PCNA-GFP+ Wrp+ midline glia (yellow arrowheads) that therefore were dividing. In **(G)** there is a notable increase in the number of Wrp+ cells. In **(E,E’)** lateral PCNA-GFP+Wrp—Dpn+ cells around the neuropile (white arrows) most likely correspond to neuropile glia. **(H)** Quantification showing phenotypic penetrance: percentage of segmentally repeated Wrp+ cell clusters that contain PCNAGFP+ cells. Fisher’s Exact test p=0.0213. **(I-L)** Over-expression of *ia-2* in glia with *repoGAL4* mildly increased the number of pH3+ mitotic cells along the midline. The pH3+ cells lacked YFP *(repoGAL4>his-YFP),* consistently with corresponding to midline glia. **(L)** Quantification in box-plots, not significantly different from controls. One Way ANOVA p=0.0995. **(A-C, E-G, I-K)** horizontal views; **(A’-C’, E’-G’, I’-K’)** transverse views. For full genotypes, sample sizes and statistical details see Table Supplement 1.

To ask whether the reprogrammed, proliferating Dpn+ cells might result in de novo neurogenesis, we first visualised cells using the *pros-*promoter, which drives expression in neural stem cells, ganglion mother cells, neurons and glia. Thus, we reasoned that this promoter would be less likely to be silenced through a cell-state transition. FlyBow was used as a reporter to visualise *pros* expressing cells. Interestingly, we could now see that the small Pros+ cells are generally neurons (Figure 9A). Over-expression of *dilp-6* with *pros-GAL4 (pros^voila^GAL4>UAS-FlyBOw, UASdilp-6)* resulted in groups of GFP+ cells (at 120h AEL) that comprised one GFP+ cell, one GFP+Dpn+ Elav— cell, and one GFP+Dpn— Elav+ cell (Figure 9A,B). These data were consistent with Dilp-6 reprogrammed glia becoming neurogenic.

**FIGURE 9.**
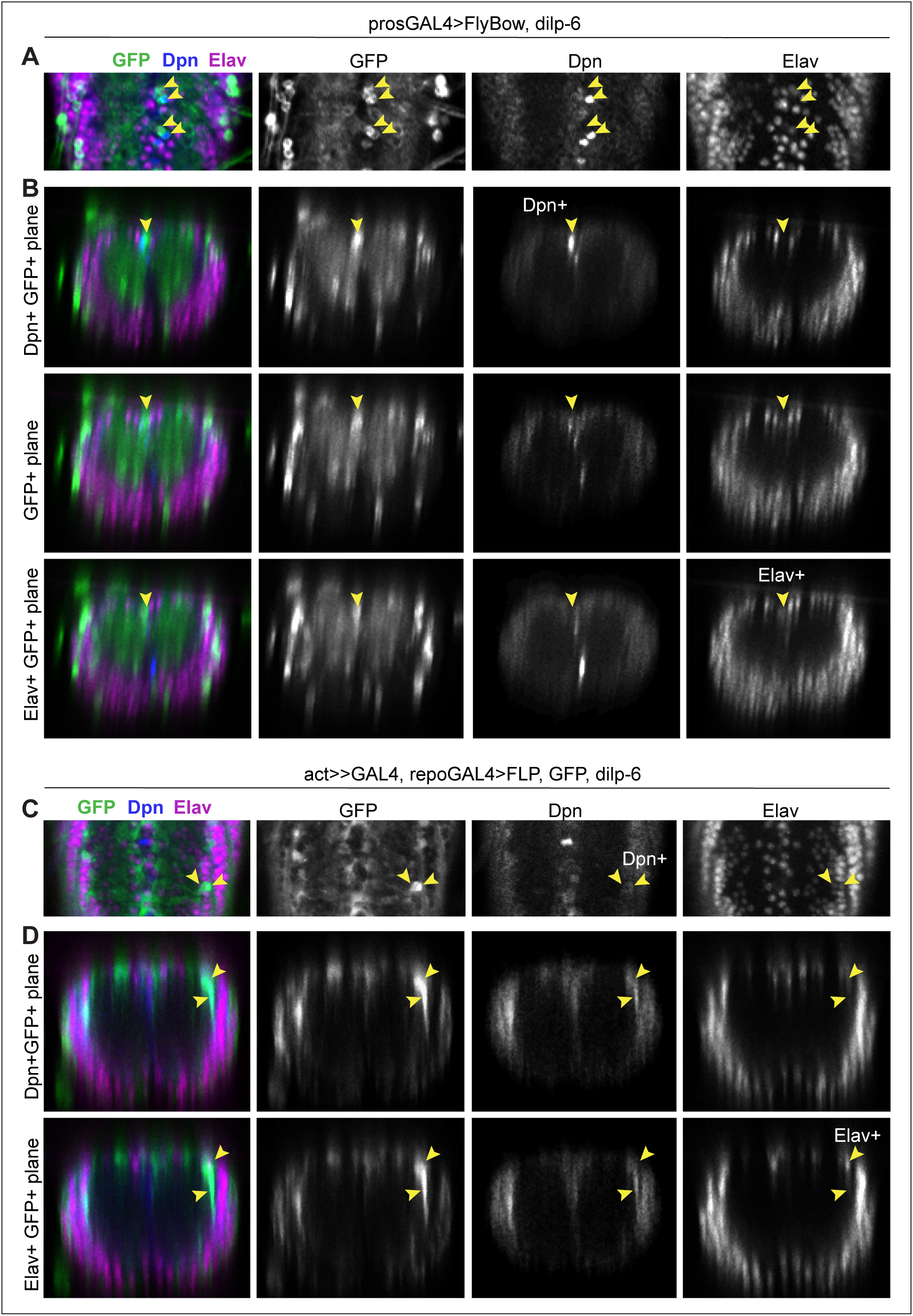
Neurons were detected from glial-derived neural stem cells. All samples were analysed at 120h AEL, after disappearance of abdominal developmental neuroblasts. **(A,B)** prosGAL4> FlyBow can reveal expression of neural stem cells, ganglion mother cells, neurons and glia. Over-expression of *dilp-6* with *prosGAL4* resulted in clusters of 3 GFP+ cells along the midline, that comprised one GFP+Dpn+ neural stem cell (top row in **B**), a GF+Dpn—Elav— progeny cell (middle row, **B**) and one GFP+Elav+ progeny neuron (bottom row, **B**). **(C,D)** Progeny cells of a glial cell-lineage were visualised with GFP, expressed originally under the control of the glial *repo* promoter, then switched using Flipase, to the permanent *actin* promoter activated only in glial cells (*actGAL4>y+>UASGFP/UAS-FLP; repoGAL4/Dilp-6).* Over-expression of *dilp-6* resulted in clusters of 2-GFP+ cells that comprised a GFP+Dpn+ neural stem cell and 2 progeny GFP+Elav+ neuronal progeny cells. **(A, C)** Horizontal and **(B,D)** transverse views. For full genotypes, sample sizes and statistical details see Table Supplement 1.

To further verify that neurons could be generated by Dilp-6 from glia, we used a second lineage-tracing method. We over-expressed *dilp-6* and *flippase (FLP)* in glia, to flip-out a stop codon placed between the *actin* promoter and GAL4, to swap the expression of the reporter GFP from being controlled by the glial *repo* promoter, to the constant *actin* promoter (*actin>y+STOP>GAL4 UASGFP/UAS-FLP; repoGAL4/dilp-6*). Thus, as reprogammed glial cells switched off the glial *repo* promoter and switched on neural stem cell gene expression, they and their progeny cells would still be visible with GFP. Larvae were analysed at 120h AEL. In this genetic background, over-expression of *dilp-6* resulted in lateral ectopic Dpn+ cells that were also GFP+ (Figure 9C,D, at 120h AEL). This showed that, like with Ia-2 and G-TRACE (Figure 7), ectopic Dpn+ cells induced by Dilp-6 originated from glia. Furthermore, there were groups of 2-3 GFP+ cells, some of which were Elav+, meaning they were neurons (Figure 9C,D). Importantly, GFP+ Elav+ cells were found near ectopic Dpn+ cells (Figure 9C,D). These data meant that glial-derived Dpn+ cells could produce neurons. We did not find any larger clusters, meaning that neurogenesis was most likely limited.

## DISCUSSION

A critical missing link to understand how to induce CNS regeneration in non-regenerating animals such as humans, had been to identify factors that interact with NG2 to induce regenerative neurogenesis. NG2 is important because NG2-glia are abundant progenitor cells present throughout life in the adult human brain, and can respond to injury (Dimou and Gotz 2014; Torper et al. 2015; Valny et al. 2017), making them the ideal cell type to manipulate to promote regeneration. However, whether NG2-glia can give rise to neurons is highly debated, and potential mechanisms remained unknown (Dimou and Gotz 2014; Vigano and Dimou 2016; Falk and Gotz 2017; Valny et al. 2017; Du et al. 2020). Here, using *Drosophila* in vivo functional genetic analysis we have identified neuronal Ia-2 as a genetic interactor of the NG2 homologue Kon, and show that it can induce a neurogenic response from glial cells via insulin signalling.

We provide evidence that Ia-2, Kon and Dilp-6 induce a regenerative neurogenic response from glia (Figure 10). In the un-injured CNS, Kon and Ia-2 are restricted to glia and neurons, respectively (Figure 10A). Ia-2 is required for neuronal Dilp-6 secretion (Cai et al. 2001; Kim et al. 2008), Dilp-6 is produced by some neurons and mostly glia, and its production depends mostly on Kon from glia. Alterations in Ia-2 levels, and increased Dilp-6, cause a non-autonomous induction of ectopic neural stem cells from glia. We do not fully understand why *ia-2* loss of function induced ectopic Dpn+. *ia-2* loss of function would cause a decrease in Dilp-6 secretion from neurons, but not from glia, as *kon* mRNA levels were unaffected (Figure 1B) and *dilp-6* expression depends mostly on glial *kon* (Figure 4C). *ia-2* loss of function caused a non-autonomous increase in Pros+ cells (Figure 2F-H), suggesting that perhaps cell-cell interactions involving Ia-2 prevent reversion to progenitor cell fate. Alternatively, as neuronal Ia-2 and glial Kon mutually exclude each other, perhaps loss of *ia-2* function might result in an increase in *kon* function that we could not detect, and with it an increase in Dilp-6 production. As Ia-2 is required for Dilp-6 secretion (Cai et al. 2001; Harashima et al. 2005; Kim et al. 2008), *ia-2* gain of function would increase Dilp-6 release triggering the Dilp-6 amplification loop (Figure 5). Conceivably, either way Dilp-6 increased and this induced Dpn. Upon injury, levels of *kon* (Losada-Perez et al. 2016) and *ia-2* expression increased (Figure 10B). Ia-2 drives secretion of Dilp-6 from neurons, Dilp-6 is received by glia, and an amplification positive feedback loop drives the further Kon and InR dependent production of Dilp-6 from cortex glia (Figure 10B). Dilp-6 can then both promote glial proliferation to generate more glia, and induce neural stem cell marker Dpn in neuropile glia – the subset known as “*Drosophila* astrocytes” and midline glia (Figure 10B,C). Ectopic Dpn+ cells were induced both upon injury and genetic manipulation of Ia-2 and Dilp-6 without injury, and originated from glia. Importantly, these glial-derived neural stem cells could divide, as revealed by the S-phase marker PCNA-GFP and the mitotic marker pH3; and could generate neurons, albeit to a rather limited extent. Altogether, Dilp-6 is relayed from neurons to cortex and then to neuropile glia. This neuron-glia communication relay could enable concerted glio- and neuro-genesis, matching interacting cell populations for regeneration (Figure 10B,D). Interestingly, Dilp-6 is also involved in non-autonomous relays between distinct CNS cell populations to activate neural stem cells and induce neuronal differentiation in development (Sousa-Nunes et al. 2011; Fernandes et al. 2017).

**FIGURE 10.**
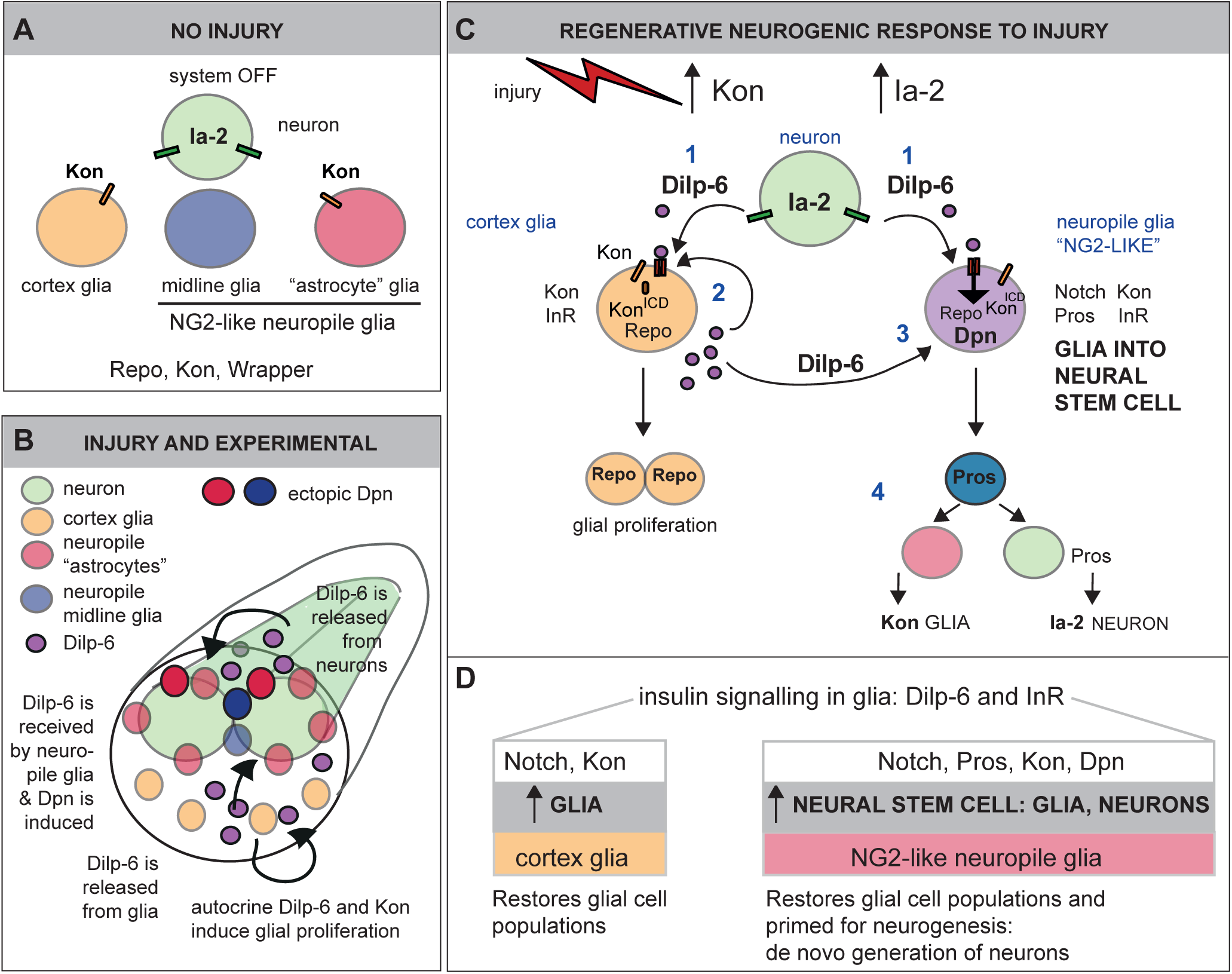
Ia-2 and Dilp-6 drive a regenerative neurogenic response to CNS injury. (A) In the abdominal larval VNC, neurons have Ia-2, glia have Kon, and Ia-2 and Kon are mutually exclusive; non-midline glia have the transcription factor Repo and midline glia the membrane protein Wrapper. In the normal, uninjured abdominal VNC, InR is in glial cells and some neurons; Ia-2 expression is constantly present in neurons; *kon* is switched off, and there are no neural stem cells (neuroblasts). **(B)** Drawing showing that Dilp-6 can be secreted from neurons, amplified and secreted by cortex glia, and received by all glial types. Dilp-6 production and secretion depend on Kon and Ia-2, which increase in injury. **(C)** Injury to the abdominal VNC provokes a dramatic surge in Ia-2 and Kon. This drives the initial secretion of Dilp-6 from neurons (1). Secreted Dilp-6 binds InR in glia, and InR signaling may facilitate cleavage and activation of Kon. Kon^ICD^ activates glial proliferation, glial cell fate gene expression and expression of *dilp-6* in cortex glia (2). In an autocrine Kon and InR dependent manner, Dilp6 sets off a positive feedback loop that amplifies Dilp-6 production from cortex glia (2). Once secreted, Dilp-6 and InR signaling cause the up-regulation of Dpn+ in neuropile glia - including Notch+ Pros+ lateral (astrocytes) and Wrp+ midline glia (3). Neuropile glia can stochastically switch on Dpn. Glial-derived Dpn+ neural stem cells can divide, and generate new neurons - although to a rather limited extent (4). After cell division, Kon may determine whether daughter cells become glia, to the exclusion of Ia-2. **(D)** Insulin signalling involving Ia-2, Dilp-6 and InR can increase cell number of various glial cell types - including cortex glia, neuropile astrocytes and midline glia - induce neural stem cells, and potentially generate new neurons. The neurogenic potential of glia may depend on the availability of Notch and Pros, and the downregulation of Kon. Together, these genes can potentially induce neurogenesis and gliogenesis, matching cell populations for regeneration.

We have demonstrated that ectopic neural stem cells originate from glia. In principle, regenerative neurogenesis could occur via direct conversion of glia into neurons, glial de-differentiation, or neuronal de-differentiation. Neuronal de-differentiation occurs both in mammals and in *Drosophila* (Froldi et al. 2015). However, in most animals, neural stem cells in the adult CNS and upon injury, are generally distinct from developmental ones, and can originate from hemocytes, but most often, glial cells (Tanaka and Ferretti 2009; Dimou and Gotz 2014; Falk and Gotz 2017; Simoes and Rhiner 2017; Du et al. 2020). In the mammalian brain, radial glia in the hippocampus respond to environmental challenge by dividing asymmetrically to produce neural progenitors that produce neurons (Shtaya et al. 2018); and astrocytes and NG2-glia can generate neurons, particularly in response to stroke, excitoxic injury, and genetic manipulations (Heinrich et al. 2014; Dimou and Gallo 2015; Peron and Berninger 2015; Du et al. 2020). Furthermore, genetic manipulation can lead to the direct conversion of NG2-glia into neurons (Torper et al. 2015; Pereira et al. 2017). Our findings that Dilp-6 and InR can induce *dpn* expression are reminiscent of their functions in the induction of neural stem cells from quiescent progenitors in development (Chell and Brand 2010; Sousa-Nunes et al. 2011; Gil-Ranedo et al. 2019). However, the Dpn+ cells induced upon injury and after development, are distinct from the developmental neural stem cells normally induced by Dilp-6 in multiple ways. Firstly, in injuries carried out in third instar larvae, the induced neural stem cells were more numerous than normal neural stem cells. Secondly, in injuries carried out late in wandering larvae, Dpn+ cells were found after normal developmental neural stem cells have been eliminated through apoptosis (Bello et al. 2003). Thirdly, Dpn+ cells were found in dorsal ectopic locations not normally occupied by developmental neural stem cells. In all injury and genetic manipulation experiments involving over-expression of either *ia-2* or *dilp-6*, ectopic Dpn+ cells were located along the midline and surrounding the neuropile, in positions normally occupied by glia. We demonstrated that ectopic Dpn+ originated from neuropile glia, including both midline glia and “Drosophila astrocytes”. Firstly, ectopic Dpn+ cells did not have Ia-2YFP, which is expressed in all neurons. Secondly, ectopic Dpn+ cells surrounding the neuropile occupied positions of astrocytes and were Repo+, and Repo— Dpn+ along the midline were Wrp+. Thirdly, the glial origin of the ectopic Dpn+ cells was demonstrated using two cell-lineage tracing methods (G-TRACE and glial activation of the *actin* promoter) whereby the expression initiated from the glia *repo* promoter was turned permanent despite cell state transitions. Thus, Ia-2 and Dilp-6 could reprogramme glial cells in vivo into neural stem cells.

Our data showed that the ectopic *ia-2* and *dilp-6* induced neural stem cells could divide and generate neurons. Dilp-6 induced glial-derived Dpn+ cells also expressed the S-phase marker PCNA-GFP, and Ia-2 also induced mitosis in midline glia. We could not detect mitotic cells surrounding the neuropile, but mitosis is brief, and could have easily been missed. The Dilp-6 induced ectopic Dpn+ cells could generate neurons that could be traced with GFP expression from their glial origin. Thus, ectopic neural stem cells induced by Dilp-6 can divide and produce neuronal progeny cells. However, the clusters of GFP+ cells originating from the in vivo reprogrammed glial cells were rather small, indicating that although neurogenesis is possible in the late larvae, it is extremely constrained. This could be due to the context of the *Drosophila* larva, where time is limited by pupariation. Injury and genetic manipulation in late larvae may not allow sufficient time for cell lineages to progress, before pupariation starts. Pupariation and metamorphosis bring in a different cellular context, which could interefere with regenerative neuronal differentiation. Alternatively, Ia-2 and Dpn may not be sufficient to carry neurogenesis through either. For instance, gain of *ia-2* function resulted only in Dpn+ but not Pros+ or Eve+ cells, suggesting that Ia-2 and Dpn are not sufficient for neuroblasts to progress to ganglion mother cells and neurons. Furthermore, ectopic Dpn+ cells still had Repo. To generate neurons, glia may not only require the expression of neural stem cell markers such as *dpn*, but also to receive other yet unknown signals (Figure 10B). In mammals, injury creates a distinct cellular environment that prompts glial cells to generate different cell types than in the un-injured CNS. For instance, elevated Sox-2 is sufficient to directly reprogramme NG2-glia into neurons, but only upon injury (Heinrich et al. 2014). Whereas during normal development NG2-glial cells may only produce oligodendrocyte lineage cells, upon injury they can also produce astrocytes and neurons (Dimou and Gallo 2015; Huang et al. 2018). This suggests that there are injury-induced cues for neuronal differentiation. In the future, it will be compelling to find out what signals could enhance neurogenesis from glial cells reprogrammed in vivo by insulin signalling.

Our work has revealed a novel molecular mechanism driving a regenerative neurogenic response from glia, involving Kon/NG2 and insulin signaling. Ia-2 induces an initial secretion of Dilp-6 from neurons, Dilp-6 is received by glia, and a positive feedback loop amplifies the Kon-dependent production of Dilp-6 by cortex glia, Dilp-6 is then relayed to neuropile glia, resulting in the in vivo reprogramming of glial cells into neural stem cells. This mechanism can induce both glial regeneration and neural stem cells from glia, potentially also neurons, matching interacting neuronal and glial cell populations. Not all neuropile glia were necessarily converted to neural stem cells, meaning the process is stochastic. Such a mechanism may also operate in mammals. In fact, Ia-2 has universal functions in dense core vesicles to release insulin (Cai et al. 2001; Harashima et al. 2005; Kim et al. 2008; Nishimura et al. 2010; Cai et al. 2011). Insulin-like growth factor 1 (IGF-1) induces the production of astrocytes, oligodendrocytes and neurons from progenitor cells in the adult brain, in response to exercise (Nieto-Estevez et al. 2016; Mir et al. 2017). The transcription factor Sox-2 that can switch astrocytes to neural stem cells and produce neurons, is a downstream effector of InR/AKT signaling (Mir et al. 2017). NG2 also interacts with downstream components of the InR signalling pathway (e.g. PI3K-Akt-mTOR) to promote cell cycle progression and regulate the expression of its downstream effectors in a positive feedback loop (Sakry et al. 2015; Nayak et al. 2018). Together, all of these findings indicate that Ia-2, NG2/Kon and insulin signaling have a common function across animals in reprogramming glial cells into becoming neural stem cells.

Intriguingly, *dpn* was only induced in neuropile associated glial cells, but not other glial types, thus perhaps only the former have neurogenic potential. Of the neuropile glia, *Drosophila* “astrocytes” and midline glia express *Notch, pros* and *kon*, as well as *InR*. The cells frequently called “astrocytes” share features with mammalian NG2-glia (Losada-Perez et al. 2016; Hidalgo and Logan 2017; Kato et al. 2018). In mammals, the combination of Notch1, Prox1 and NG2 is unique to NG2-glia, and is absent from astrocytes (Cahoy et al. 2008). Perhaps Ia-2 and Dilp-6 can only induce neural stem cells from NG2-like glia bearing this combination of factors. Notch activates glial proliferation and *kon* expression in *Drosophila* (Losada-Perez et al. 2016), and in the mammalian CNS, Notch promotes NG2-glia proliferation and maintains the progenitor state, whereas its downregulation is required to induce both glial and neuronal differentiation (Yamamoto et al. 2001; Ables et al. 2010; Piccin et al. 2013; Falk and Gotz 2017). In *Drosophila,* Notch and Pros also regulate *dpn* expression: Notch activates *dpn* expression promoting stemness, and Pros inhibits it, promoting transition to ganglion mother cell and neuron (Vaessin et al. 1991; San-Juan and Baonza 2011; Babaoglan et al. 2013; Bi and Kuang 2015). Thus, only glial cells with Notch and Pros may be poised to modulate stemness and neuronal differentiation. We showed that *InR* is expressed in neuropile glia, and insulin signaling represses FoxO, which represses *dpn,* thus ultimately activates *dpn* expression (Siegrist et al. 2010). As Notch and insulin signaling positively regulate *dpn* expression (Vaessin et al. 1991; Siegrist et al. 2010; San-Juan and Baonza 2011; Babaoglan et al. 2013; Bi and Kuang 2015), and injury induces a Notch-dependent up-regulation of Kon (Losada-Perez et al. 2016), which activates *dilp-6* expression, and of Ia-2, which secretes Dilp-6, our data indicate that Notch-Kon/NG2-insulin synergy triggers the activation of *dpn* expression. Importantly, we found no evidence that Kon functions in neural stem cells. Thus, perhaps induced neural stem cells can generate only glia from daughter cells that inherit Kon, on which Repo and glial cell fate depend, or generate neurons, from daughter cells that lack Kon, but have Pros, on which Ia-2 depends (Figure 10B). Thus, upon injury, Notch, Pros, Kon/NG2, Ia-2 and insulin signalling function together to enable the regenerative production of both glial cells, and neural stem cells from glia (Figure 10B,C).

To conclude, a neuron-glia communication relay involving Ia-2, Dilp-6, Kon and InR is responsible for the induction of the neural stem cells from glia, their proliferation and limited neurogenesis. Neuronal Ia-2 and Dilp-6 trigger two distinct responses in glia: 1) in cortex glial cells, insulin signaling boosts Kon-dependent amplification of Dilp-6, glial proliferation and glial regeneration; 2) in neuropile-associated NG2-like glial cells, Ia-2 and Dilp-6 also unlock a neurogenic response, inducing neural stem cell fate. As a result, these genes can drive the production of both glial cells and neurons after injury, enabling the matching of interacting cell populations, which is essential for regeneration.

## MATERIALS AND METHODS

### Fly stocks and genetics

Fly stocks used are listed in Table 1 below. Stocks carrying combinations of over-expression and RNAi, or RNAi and mutants, etc., were generated by conventional genetics. N^ts^ mutants were raised at 18°C to enable normal embryogenesis, and switched to 25°C from larval hatching to the third instar larval wandering stage to cause N loss of function. For all experiments, larvae bearing balancer chromosomes were identified by either using the fluorescent balancers CyO Dfd-YFP and TM6B Dfd-YFP or using the balancer SM6a-TM6B Tb^—^, which balances both the second and third chromosomes, and discarded. For the genetic screens, larvae with fluoresencent VNCs (i.e. repoGAL4>UAS-FlyBow or elavGAL4>UAS-FlyBow) were selected.

### Crush injury in the larval VNC

Crush injury in the larval CNS was carried out as previously reported (Losada-Perez et al. 2016), and only lesions in the abdominal VNC were analysed. Larval collections were staged by putting the G0 flies in an egg laying chamber for 2h, then collecting the F1 larve some time later, as indicated next. Larvae were placed on a chilled petri-dish with agar over ice. Crush injury was carried out by pinching with fine forceps the GFP-bearing VNCs under UV light using a fluorescence dissecting microscope: (1) at 74-76h after egg laying (AEL); VNCs were then left to carry on developing at 25°C, and were dissected either 5-7h or 24h post-injury (PI); (2) at 96 h, kept at 25°C and dissected and fixed 6h PI; (3) at 105h AEL, kept at 25°C and dissected 24h PI; (4) at 117h AEL, kept at 25°C, and dissected 12h PI. Dissected and fixed VNC were then processed for antibody stainings following standard procedures.

### Molecular cloning

The UAS-konICD-HA construct was generated from EST LD31354 via PCR amplification with Kappa HiFi PCR kit (Peqlab) and subsequent cloning using the Gateway cloning system (Invitrogen) according to manufacturers instructions. Primers used were kon^ICD^ fwd comprising the CACC-sequence at the 5’-end (<u>CCAC</u>AGGAAACTGAGAAAGCACAAGGC) for direct cloning of the PCR product (482 bp) into the entry vector pENTR/D-Topo, and kon^ICD^ rev (AAACC<u>TTA</u>CACCCAATACTGATTCC) including the endogenous stop-codon, underlined. Destination vector was pTHW for tagging the ICD on the N-terminus with HA, including a 5xUAS cassette and P-element ends for transformation. These destination vectors were developed by the Murphy-Lab at Carnegie Institution of Science, Baltimore, MD, USA, and can be obtained from the *Drosophila* Genomics Resource Center at Indiana University, USA. Transformant fly strains were generated by BestGene Inc, Chino Hills, CA, USA following a standard tranposase-mediated germline transformation protocol.

A UAS-ia-2 construct was generated using Gateway cloning (Invitrogen, as above). Ia-2 cDNA was generated by reverse-transcription PCR of purified mRNA from Oregon R flies, and cloned into pDONR. Subsequently, a standard PCR amplification was performed using Phusion High-Fidelity (Fisher Scientific), primers Ia-2F (5’-ATGGCACGCAATGTACAACAACGGC) and ia-2-stopR (5’ -CTTCTTCGCCTGCTTCGCCGATTTG), and the resulting PCR product (3918bp) was cloned into pGEM®-T Easy Vector (Promega). Subsequently, a Phusion High Fidelity PCR amplification was carried out using Gateway primers Ia-2attB F1 (5’ -ggggacaagtttgtacaaaaaagcaggcttcATGGCACGCAATGTACAACAACGGC) and Ia-2attB R1 (5’ –ggggaccactttgtacaagaaagctgggtcCTTCTTCGCCTGCTTCGCCGATTTG), and plasmid pGEM-ia-2 as template. Using Gateway cloning, the PCR product (3979bp) was cloned first into pDONR^221^ and subsequently into the pUAS-gw-attB destination vector, for *ϕ*C31 transgenesis. The construct was injected by BestGene Inc. to generate transgenic flies bearing UAS-ia-2 at the attP2 landing site.

### Quantitative real time reverse transcription PCR (qRT-PCR)

qRT-PCR was preformed according to standard methods and as previously described (Losada-Perez et al. 2016), with the following alteration. For each sample, 10 third instar larvae were used per genotype per replicate. At least three independent biological replicates were performed for all experiments other than in Figure 1-Figure supplement 3 A and B where two replicates were carried out on all candidates and those of interest where taken forward to carry out two further replicates. For a list of the primers used in this study please see Key Resources Table below.

**Immunostainings** were carried out following standard procedures. The following primary antibodies were used: mouse anti-Repo (1:100, DSHB); guinea pig anti-Repo (1:1000, Ben Altenhein); rat anti-Elav (1:250, DSHB); mouse anti-FasII ID4 (1:500, DSHB); mouse anti-Prospero (1:250, DSHB); guinea pig anti-Dpn (1:1000, gift of J. Skeath); mouse anti-Eve 3C10 (1:20, DSHB); rabbit anti-phospho-histone-H3 (1:250); rabbit anti-HA (1:1600, Cell Signalling Technology); rabbit anti-GFP at 1:250 (Molecular Probes); mouse anti-Wrapper (1:5, DSHB). Secondary antibodies were Alexa conjugated: Donkey anti-rabbit 488 (1:250, Molecular Probes), goat anti-rabbit 488 (1:250, Molecular Probes), goat anti-rabbit 647 (1:250, Molecular Probes), goat anti-mouse 488 (1:250, Molecular Probes), goat anti-mouse 546 (1:250, Molecular Probes), goat anti-mouse 647 (1:250, Molecular Probes), goat anti-rat 546 (1:250, Molecular Probes), goat anti-guinea pig 488 (1:250, Molecular Probes), goat anti-guinea pig 633 (1:250, Molecular Probes), goat anti-rat 647 and 660 (1:250, Molecular Probes).

### Microscopy and imaging

Image data were acquired using a Zeiss LSM710 laser scanning confocal microscope, with 25x lens and 1.00 zoom, an Olympus Fluoview FV1000, 20x lens, and a Leica SP8 laser scanning confocal microscope, with a 20x lens, 1.25 zoom. All images were taken with resolution 512×512 or 1024×1024, step 0.96μm and 1-3x averaging for all samples except for cell counting with DeadEasy that have no averaging.

Images were analysed using ImageJ. Images of horizontal sections are projections from the stacks of confocal images that span the thickness of the entire VNC, using ImageJ. Transverse views were generated using the Reslice option. Images were processed using Adobe Creative Suite 6 Photoshop and compiled with Adobe Illustrator.

### Automatic cell counting

Glial cells labelled either with anti-Repo or with repoGAL4>UAShistone-YFP were counted automatically in 3D across the thickness of the VNC using DeadEasy Larval Glia software, as previously described. Prospero+ and Dpn+ cells were counted manually in 3D (i.e. not in projections), as the signal was noisy for DeadEasy.

### Statistical analysis

Statistical analysis was carried out using Graphpad Prism. All data in this work are continuous, except for the PCNAGFP data in Figure 8H which are categorical. The latter were analysed with a non-parametric Fisher’s exact test. For all other data, tests to determine whether data were distributed normally and variances were equal were initially carried out, and thereafter if so, parametric One Way ANOVA tests where carried out when comparing more than two sample types group. Multiple comparison corrections were carried out with post-hoc Dunnett tests comparisons to set controls, or Bonferroni comparisons of all samples against all. Box plots were used to represent the distribution of continuous data, where the line within the box represents the median of the data distribution, the box comprises the 25 percentiles above and below the median, and the whiskers the lowest and highest 25 percentiles.

## ACKNOWLEDGEMENTS

We thank our labs and C. Rezaval for discussions and comments on the manuscript; S. Corneliussen, T. Schunke and S. Dietz for technical help; Y. Fan, A. Gould, Y. Jan, Y. Nung, J. Skeath and F. Schnorrer for reagents; A. Di Maio and Birmingham Advanced Light Microscopy for assistance; Bloomington Drosophila Stock Centre for fruit-flies and Developmental Studies Hybridoma Bank, Iowa for antibodies.

---

**Figure 1 – Figure Supplement 1.**
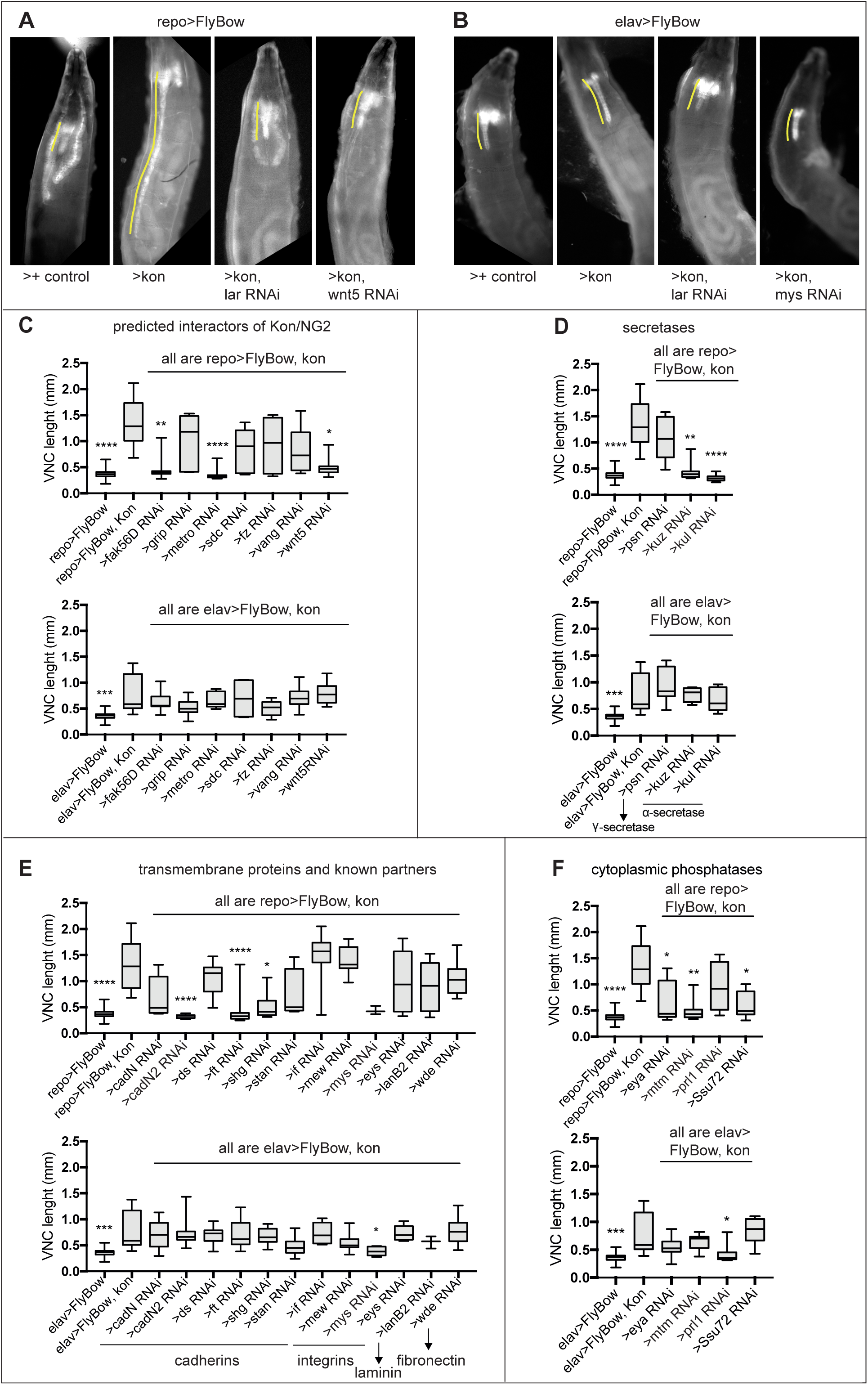
Modifier genetic screens identify genes interacting with *kon*. **(A,B)** Over-expression of *kon* in glia with *repoGAL4 (repo>UASFlyBow, UASkon-full-length)* caused a phenotype of very long ventral nerve cord **(A),** and in neurons with *elavGAL4* too, although to a lesser extent **(B).** These phenotypes were quantified by using the reporter *UASFlyBow*, and the VNC measured using ImageJ tools. RNAi knock-down of candidate genes could rescue these phenotypes, some examples are shown here. **(C-F)** The *kon* gain of function (GOF) phenotype resulting from over-expressing *kon-full-length* in either neurons or glia, could be rescued by RNAi knock-down of: **(C)** predicted interactors of Kon or NG2, most prominently in glia; Kruskal Wallis ANOVA p<0.0001, post-hoc Dunn test to *>FlyBow, kon controls.* N=4-24 VNCs. **(D)** *α* and *γ*-secretases that cleave NG2 and Notch, from glia; Kruskal Wallis ANOVA p<0.0001, post-hoc Dunn test to *>FlyBow, kon controls.* N=4-24 VNCs. **(E)** Known Kon partners, e.g. integrins, and other transmembrane proteins from neurons; Kruskal Wallis ANOVA p<0.0001, post-hoc Dunn test to *>FlyBow, kon controls.*N=3-24 VNCs. **(F)** cytoplasmic phosphatases, from either glia or neurons; Kruskal Wallis ANOVA p<0.0001, post-hoc Dunn test to *>FlyBow, kon controls.* N=7-24 VNCs. VNC length indicated in yellow in (A,B). *Asterisks indicate multiple comparison post-hoc tests to controls: *p<0.05, **p<0.01, ***p<0.001, ****p<0.0001. For full genotypes and further statistical analysis details see Table Supplement 1.

**Figure 1 – Figure Supplement 2.**
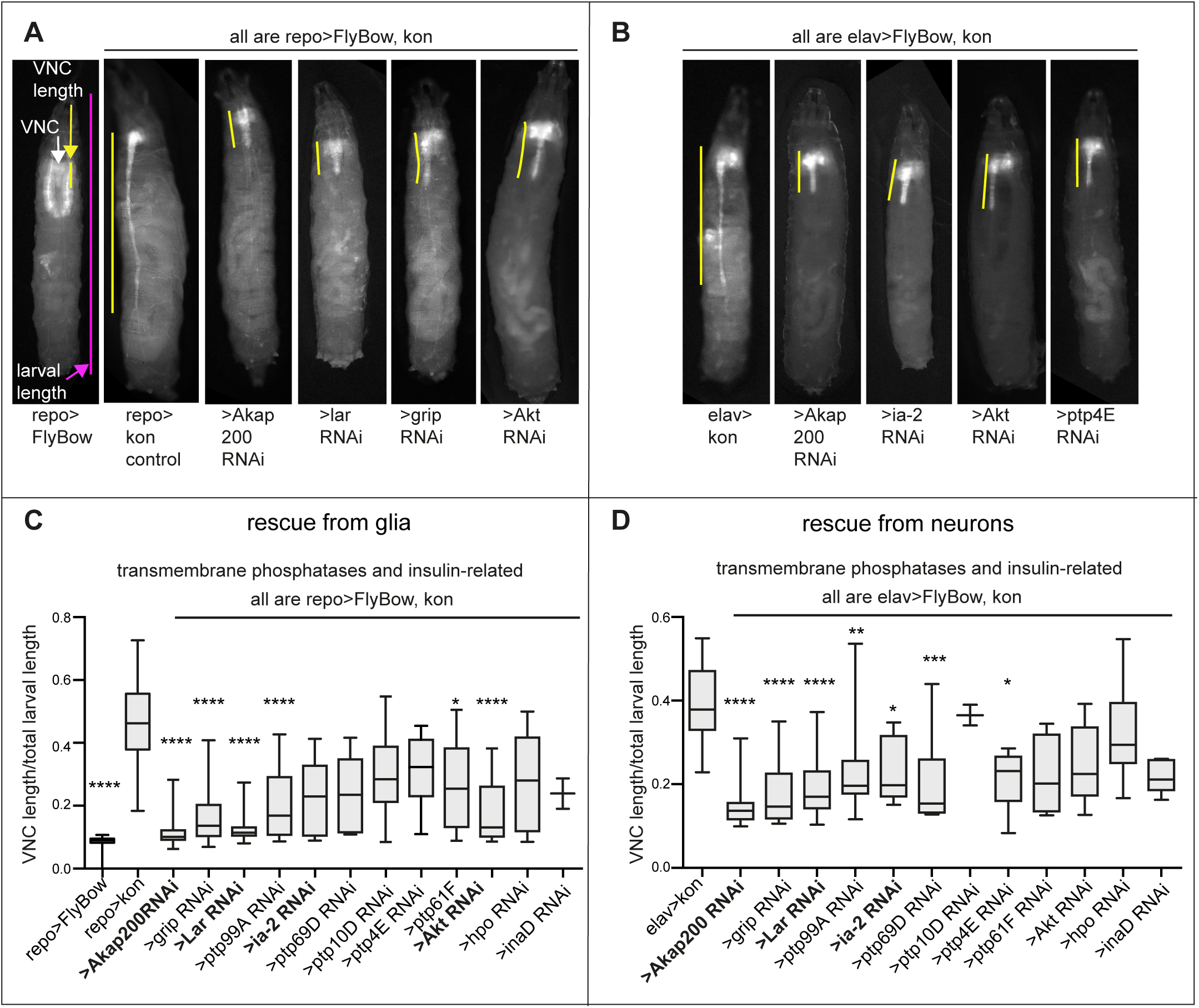
Modifier candidate genetic screens identify genes encoding transmembrane phosphatases and insulin signaling factors as interacting with *kon*. (A,B) Over-expression of *kon* in glia causes a very long VNC **(A),** and in neurons too, but to a lesser extent **(B).** RNAi knock-down of candidate genes could rescue these gain of function phenotypes, some examples are given. **(C,D)** Quantification of normalized VNC length shows rescue prominently by knock-down of most transmembrane phosphatases, the Notch-related *Akap200*, and genes functionally related to the insulin signaling pathway (*Akt, lar* and *ia-2)*, most prominently *lar.* Normalised measurements are given as a ratio of the VNC over total larval length. Kruskal-Wallis ANOVA p<0.0001, post-hoc Dunn’s test comparison to controls *repo>kon* or *elav>kon*. **(C)** N=2-28; **(D)** N=2-31. *Asterisks indicate multiple comparison post-hoc tests to controls: *p<0.05, **p<0.01, ***p<0.001, ****p<0.0001. For full genotypes and further statistical analysis details see Table Supplement 1.

**Figure 1 – Figure Supplement 3.**
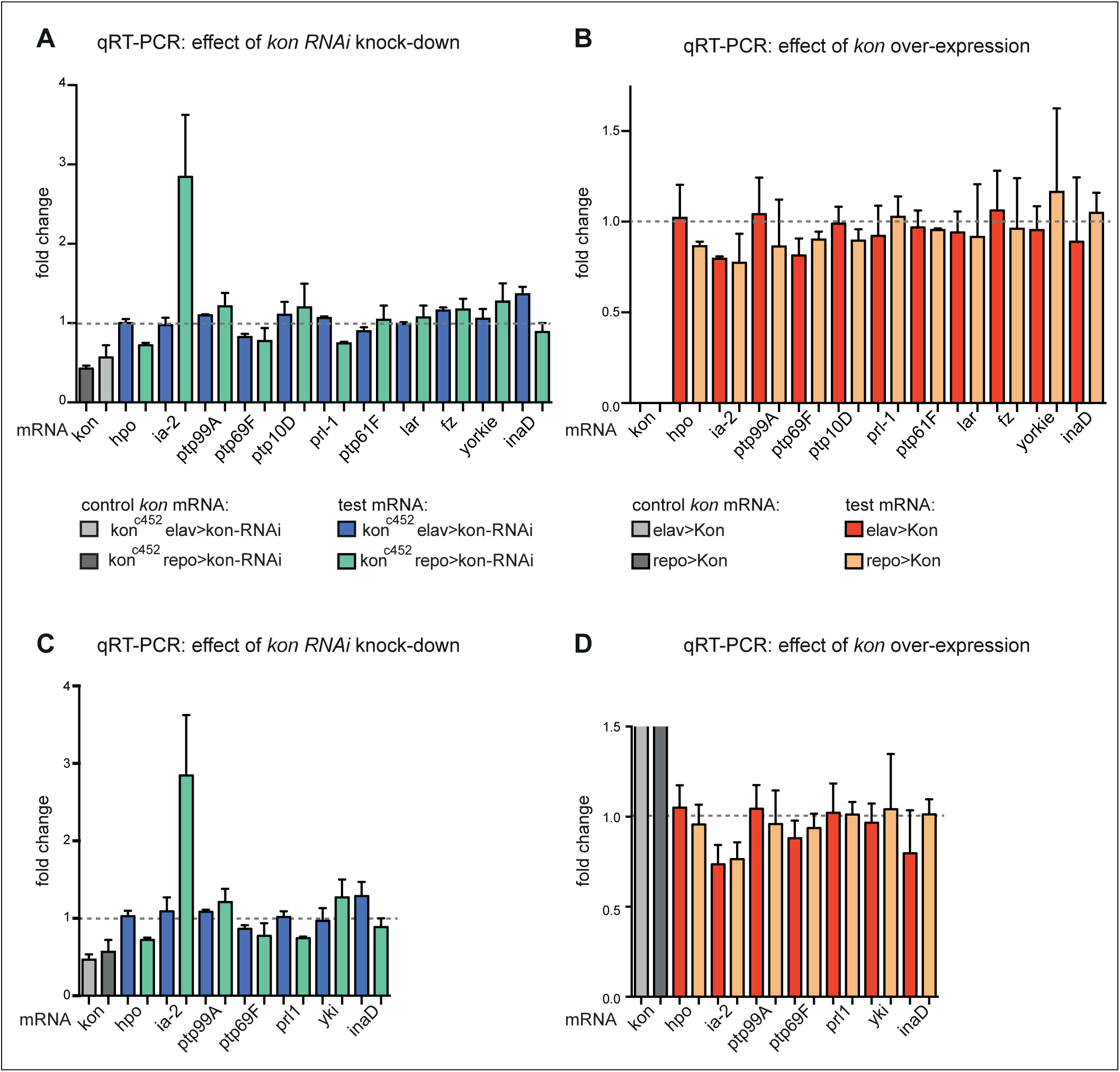
Loss and gain of *kon* function prominently affected *ia-2* expression. **(A,B)** Exploratory quantitative real-time PCR (qRT-PCR), N=2 replicates each: **(A)** showing the change in mRNA levels for candidate genes upon *kon* RNAi targeted to either neurons (with *elavGAL4*) or glia (with *repoGAL4*). *ia-2* mRNA levels increased at least 3 fold when *kon* was knocked-down in glia; **(B)** showing the effect of *kon* gain of function. *kon* over-expression in either neurons or glia decreased *ia-2* mRNA levels. The first two columns have been left cut out as they are controls with the increase in *kon* mRNA with *kon* over-expression, which are very high compared to the rest. **(C,D)** Further replicates were carried out for a selected group of genes, and they validate that *kon* prominently regulates *ia-2* expression. N=4 replicates each. **(D)** The first columns represent the very high increase in *kon* mRNA with *kon* over-expression, and they have been cut as they go well beyond this scale compared to the rest. For full genotypes and further statistical analysis details see Table Supplement 1.

**Figure 2 – Figure Supplement 1.**
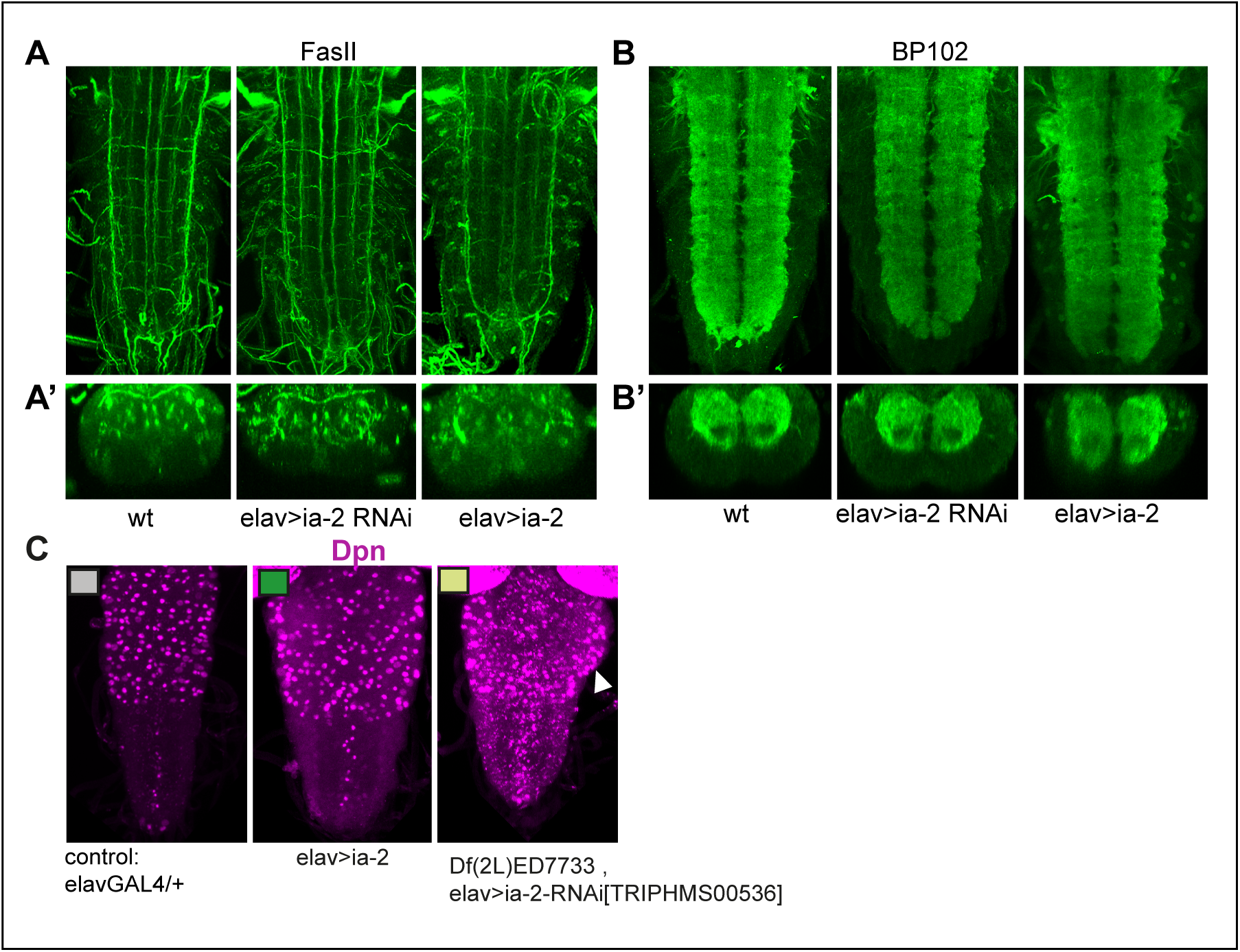
Alterations in *ia-2* levels cause no obvious neuronal phenotypes, but can induce overgrowths. (A) Neurons and their axonal fascicles are visualised with anti-FasII. N=7-11 VNCs. **(B)** Neurons and their dendrites are visualized with anti-BP102. N=9-10 VNCs. No abnormal phenotypes were observed. **(A,B)** Horizontal views; **(A’,B’)** transverse views. **(C)** Anti-Dpn visualised in larvae at 120h AEL, after disappearance of developmental neuroblasts. Loss of *ia-2* function in neurons caused over-growths in thorax (arrowhead). N=7-12 VNCs.

**Figure 5 Figure Supplement 1.**
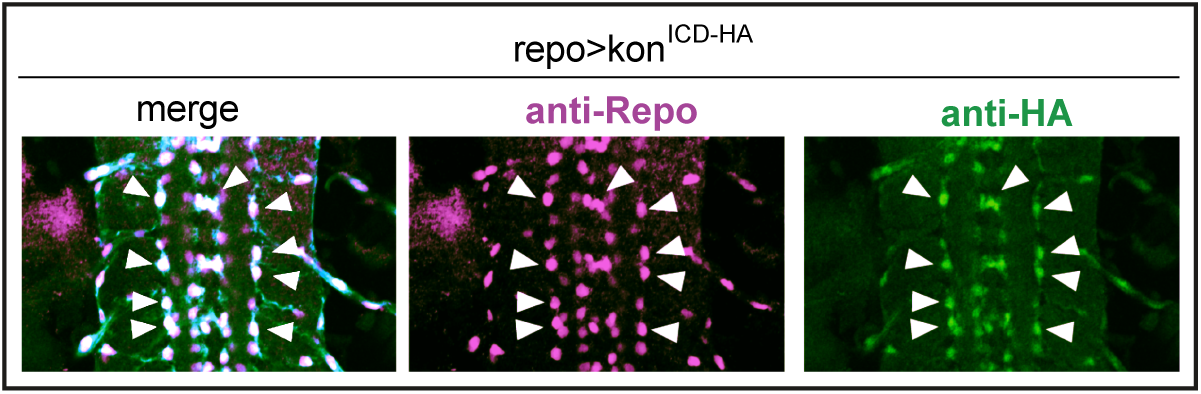
Over-expressed HA-tagged Kon^ICD^ in glia (*repoGAL4>UASkon^ICd^::HA*) visualized with anti-HA antibodies in stage 16 embryos localizes to glial nuclei stained with the pan-glial marker anti-Repo (arrows).

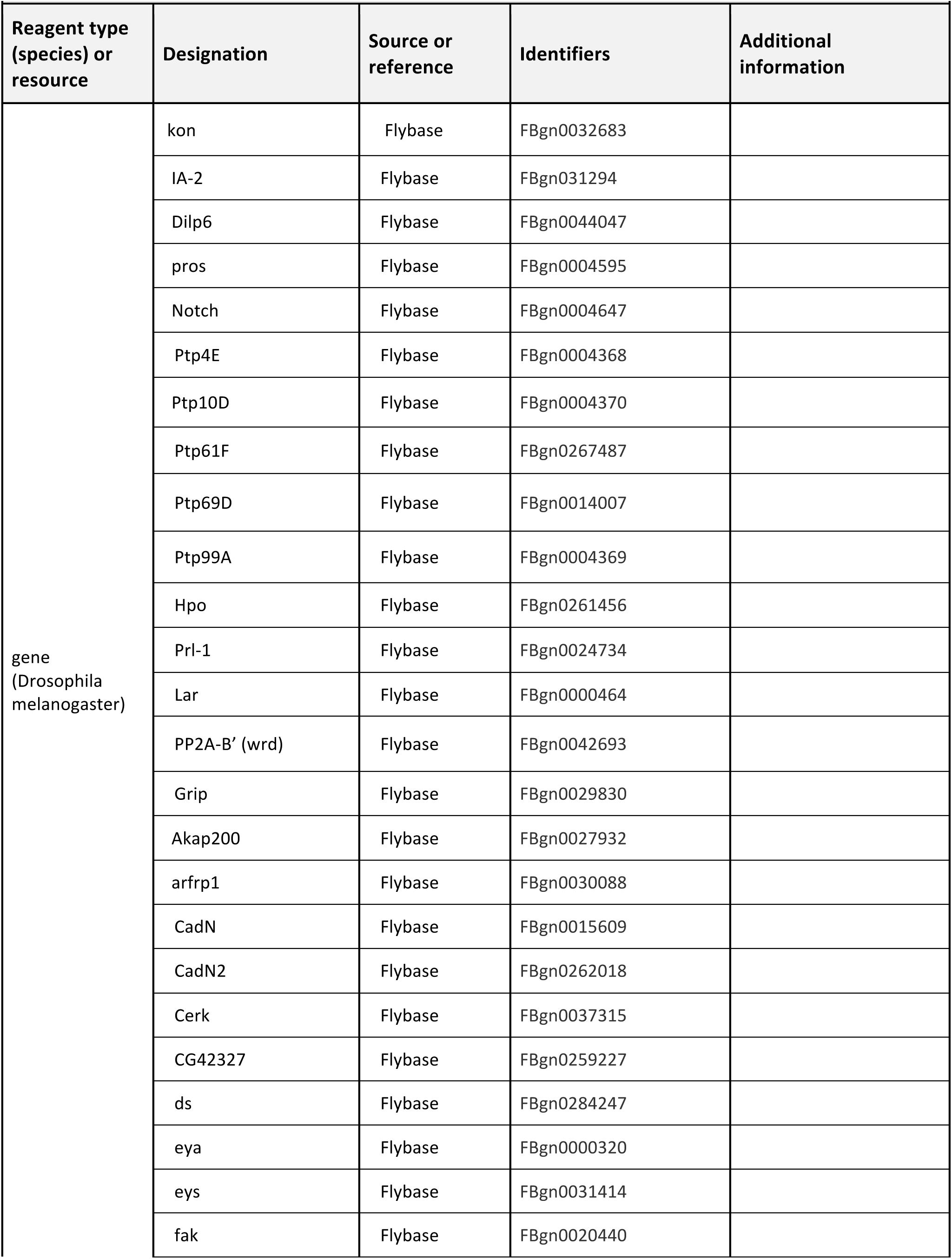
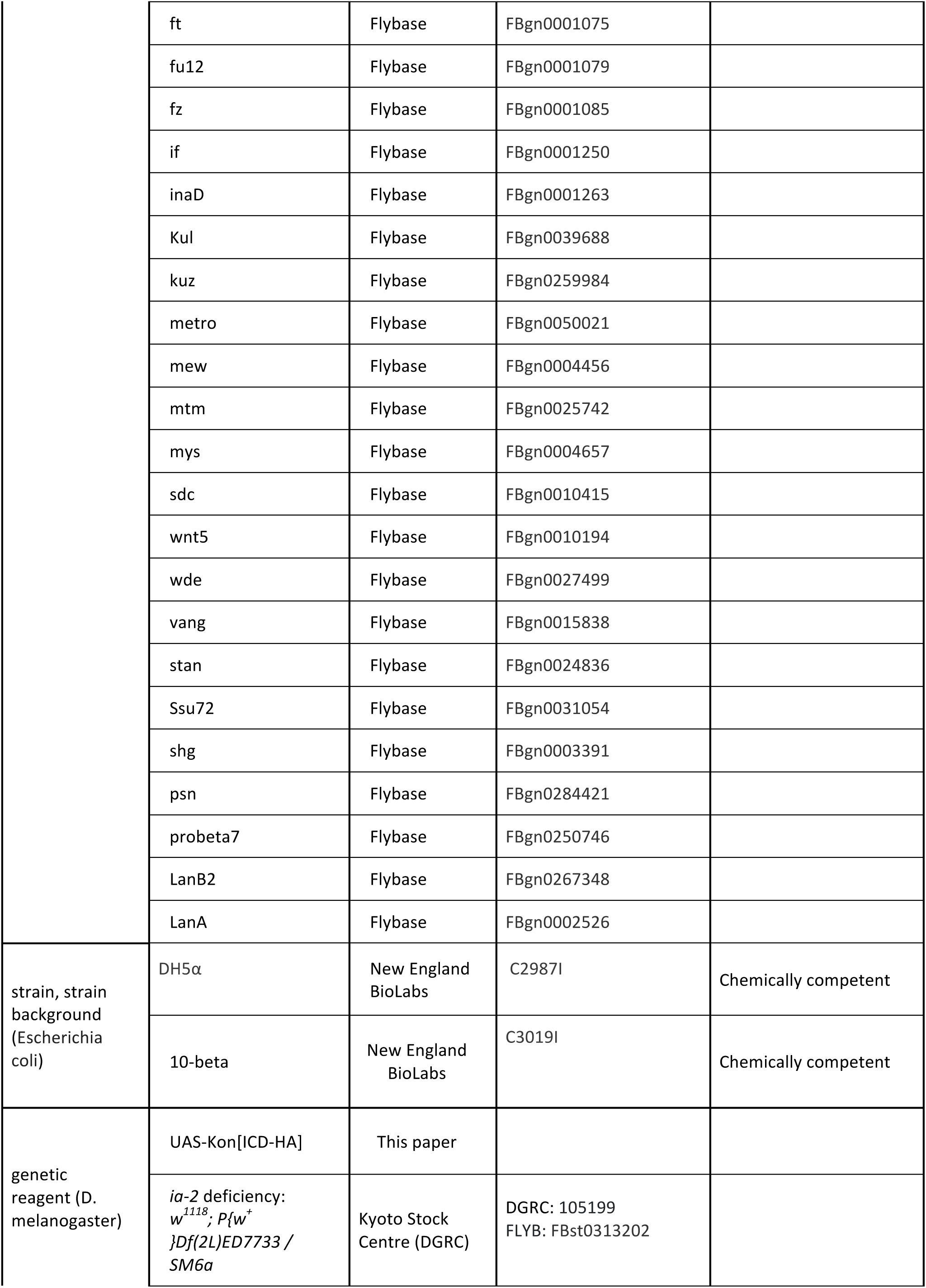
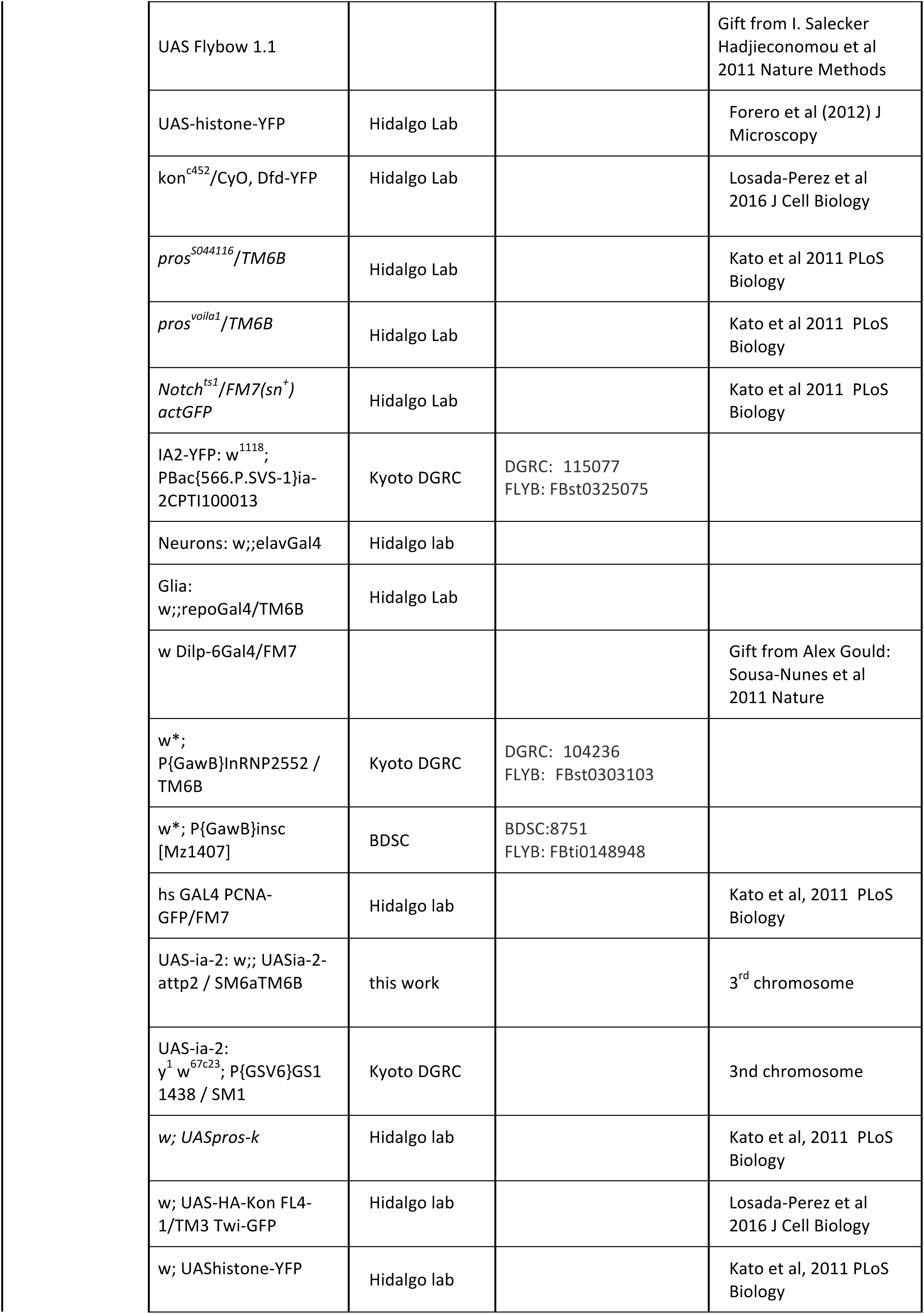
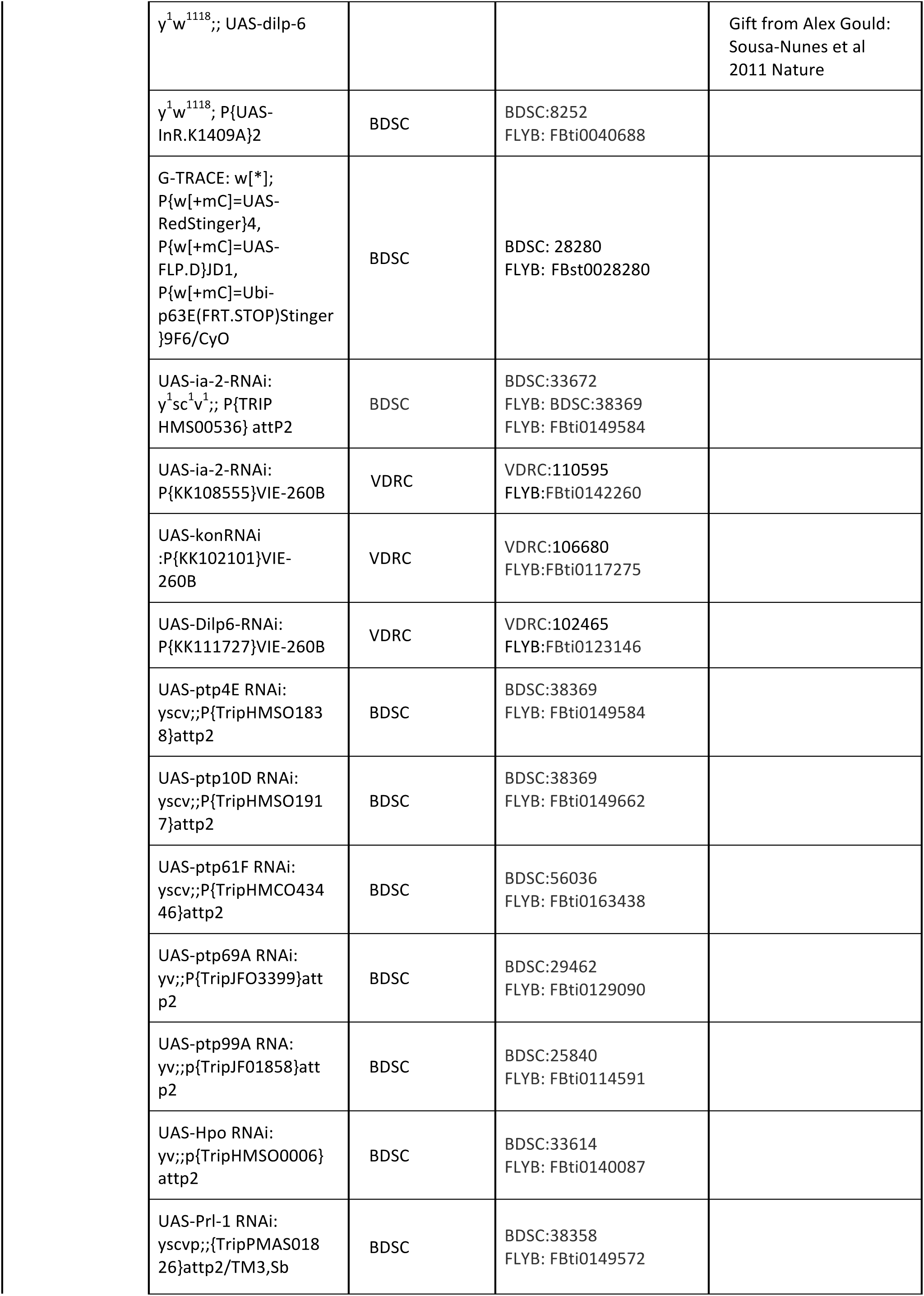
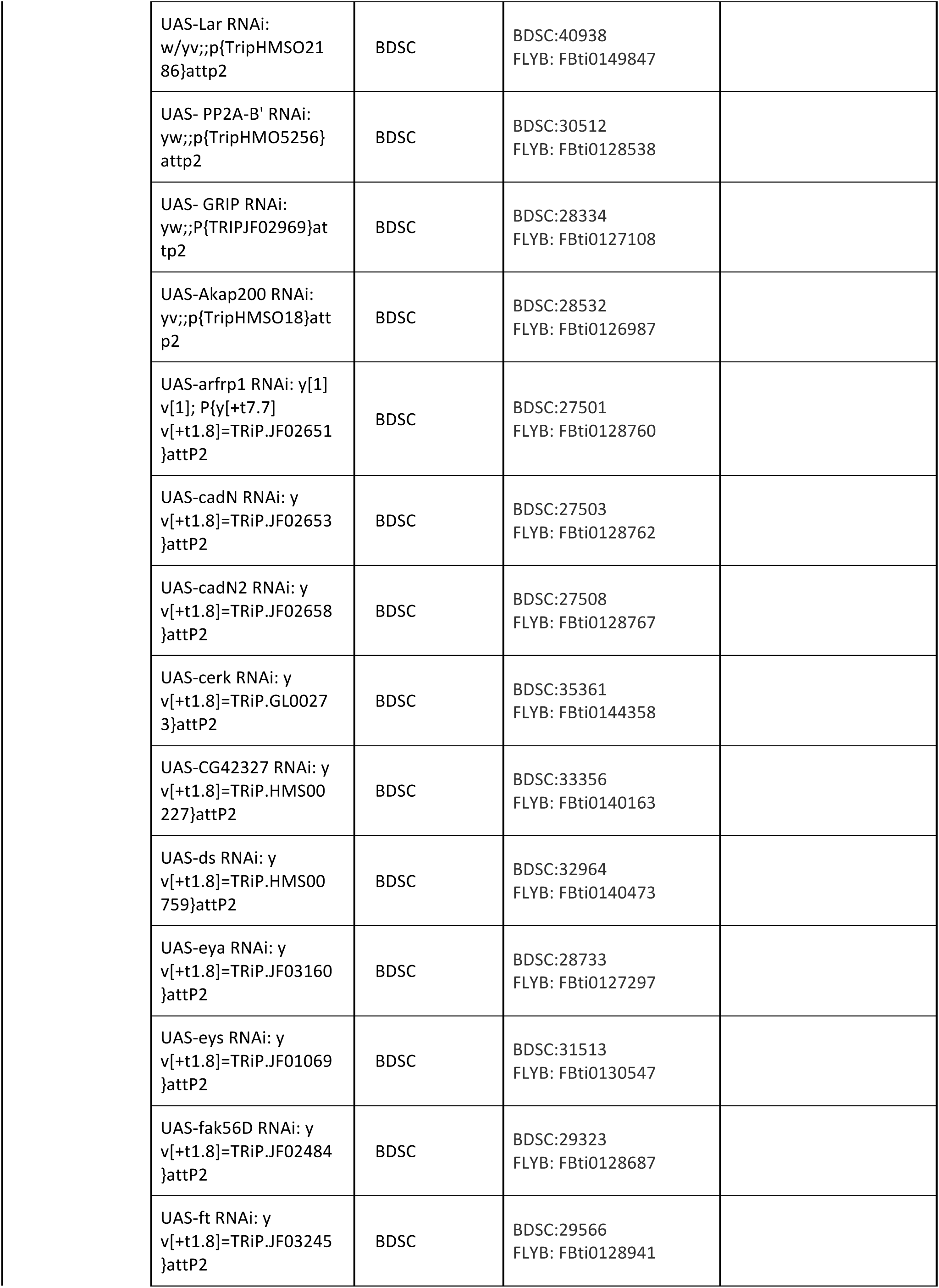
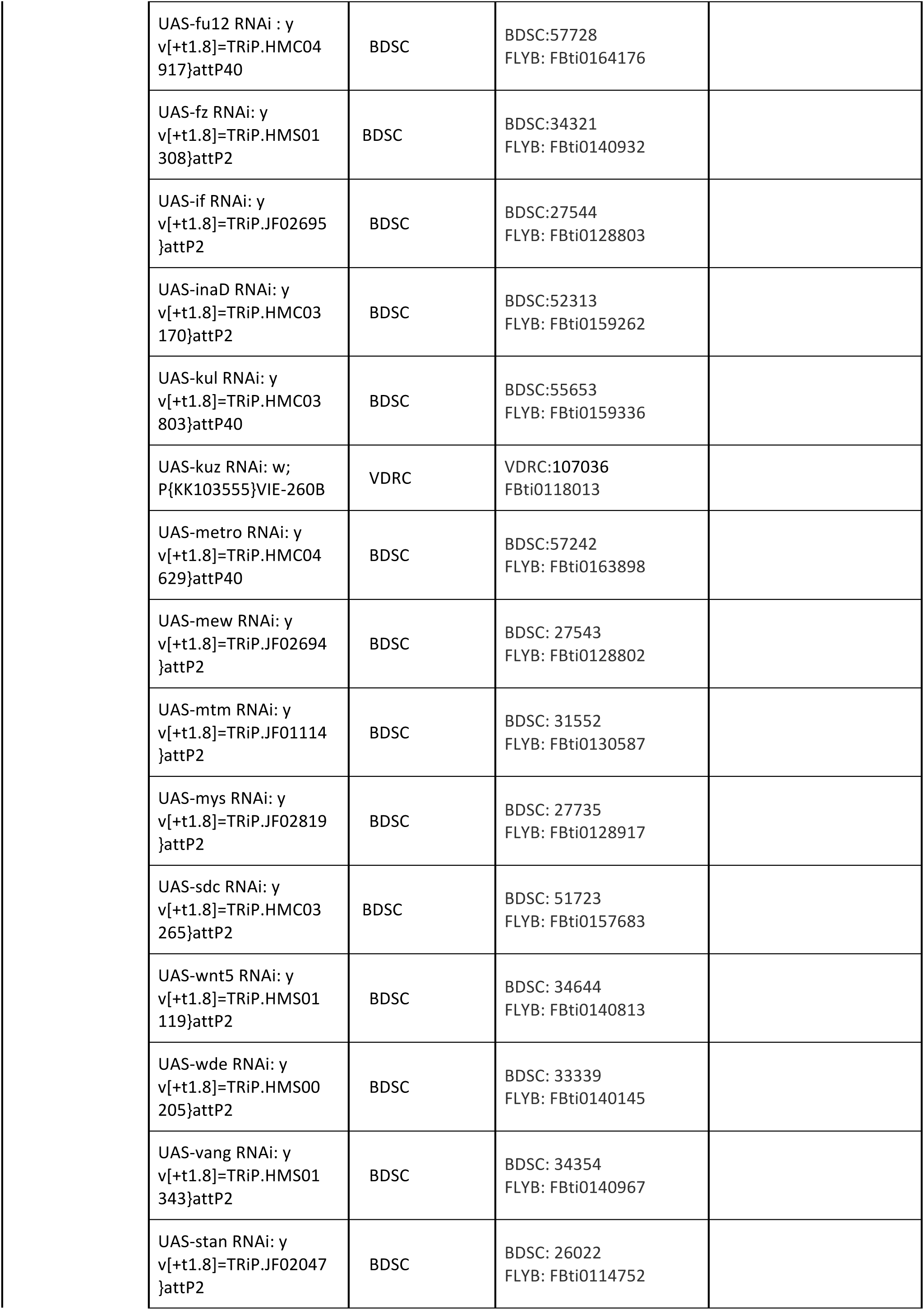
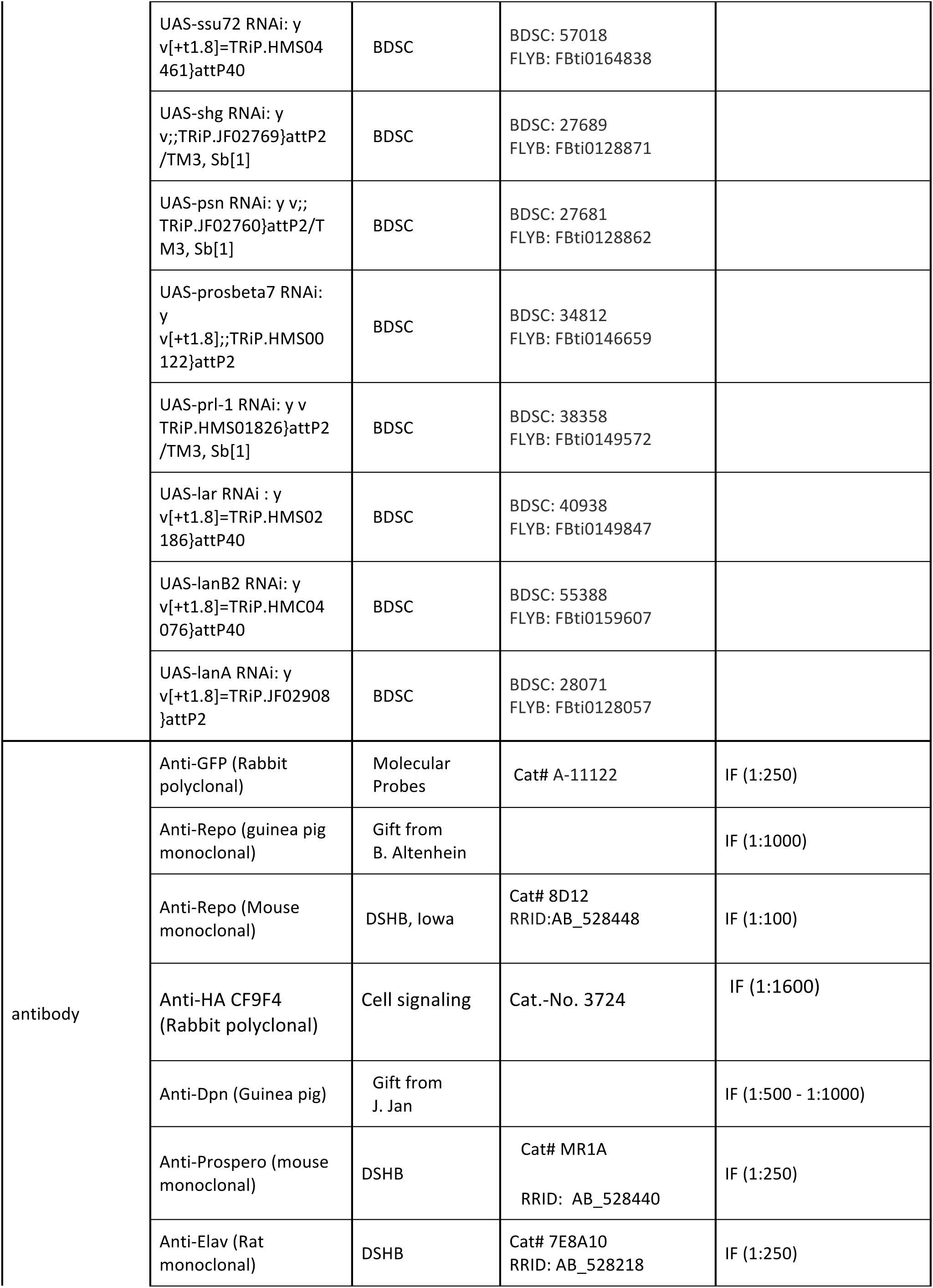
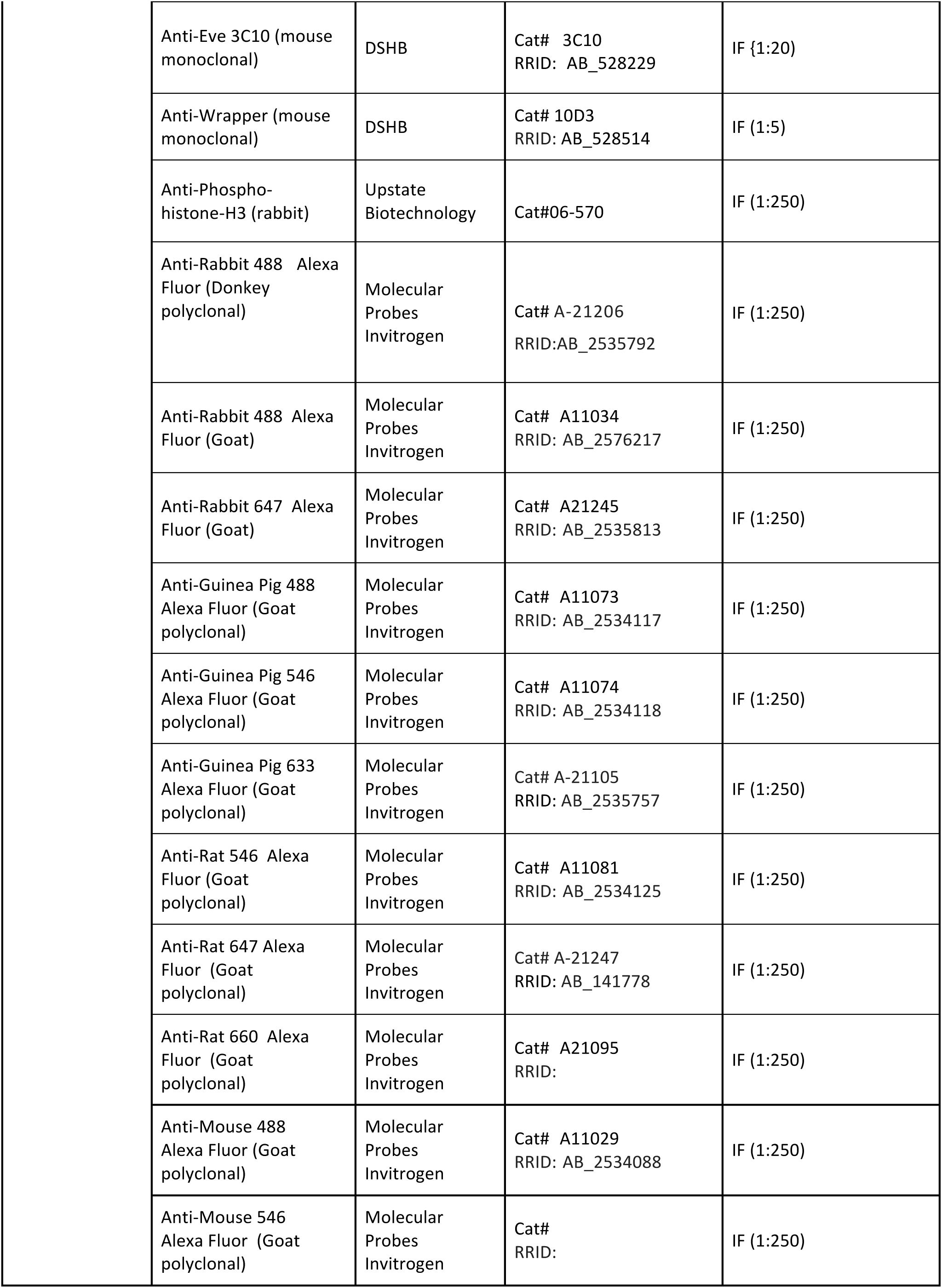
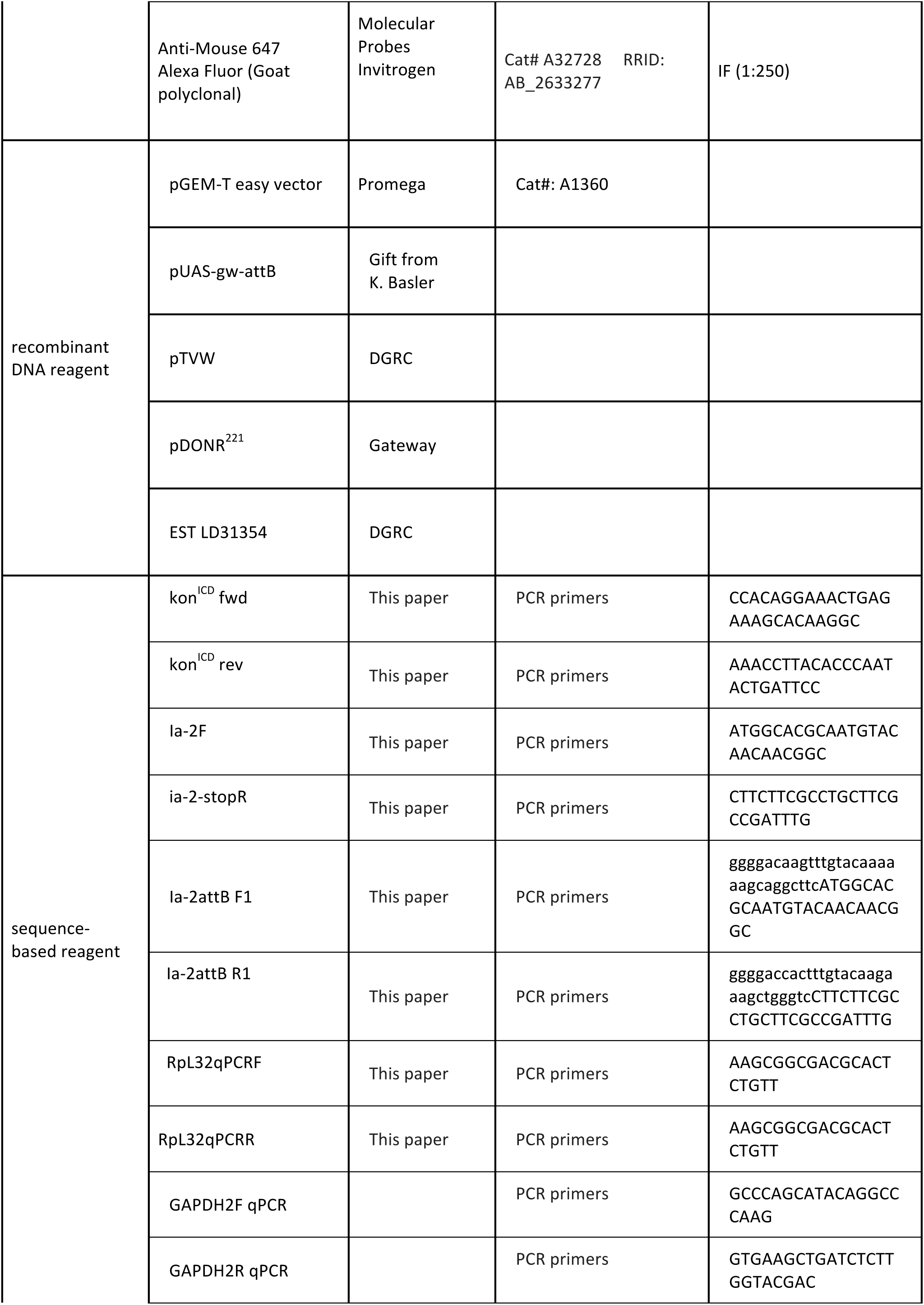
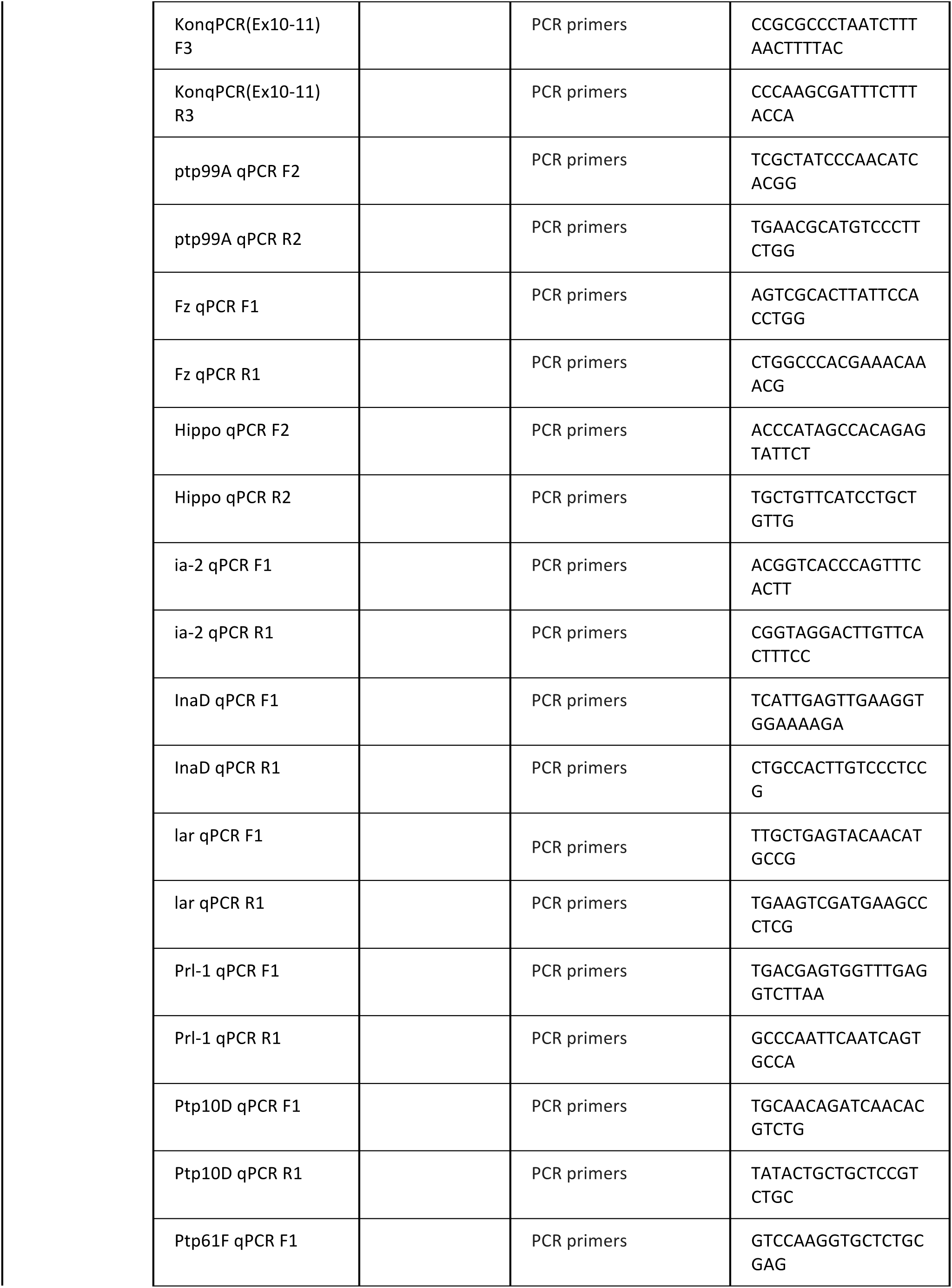
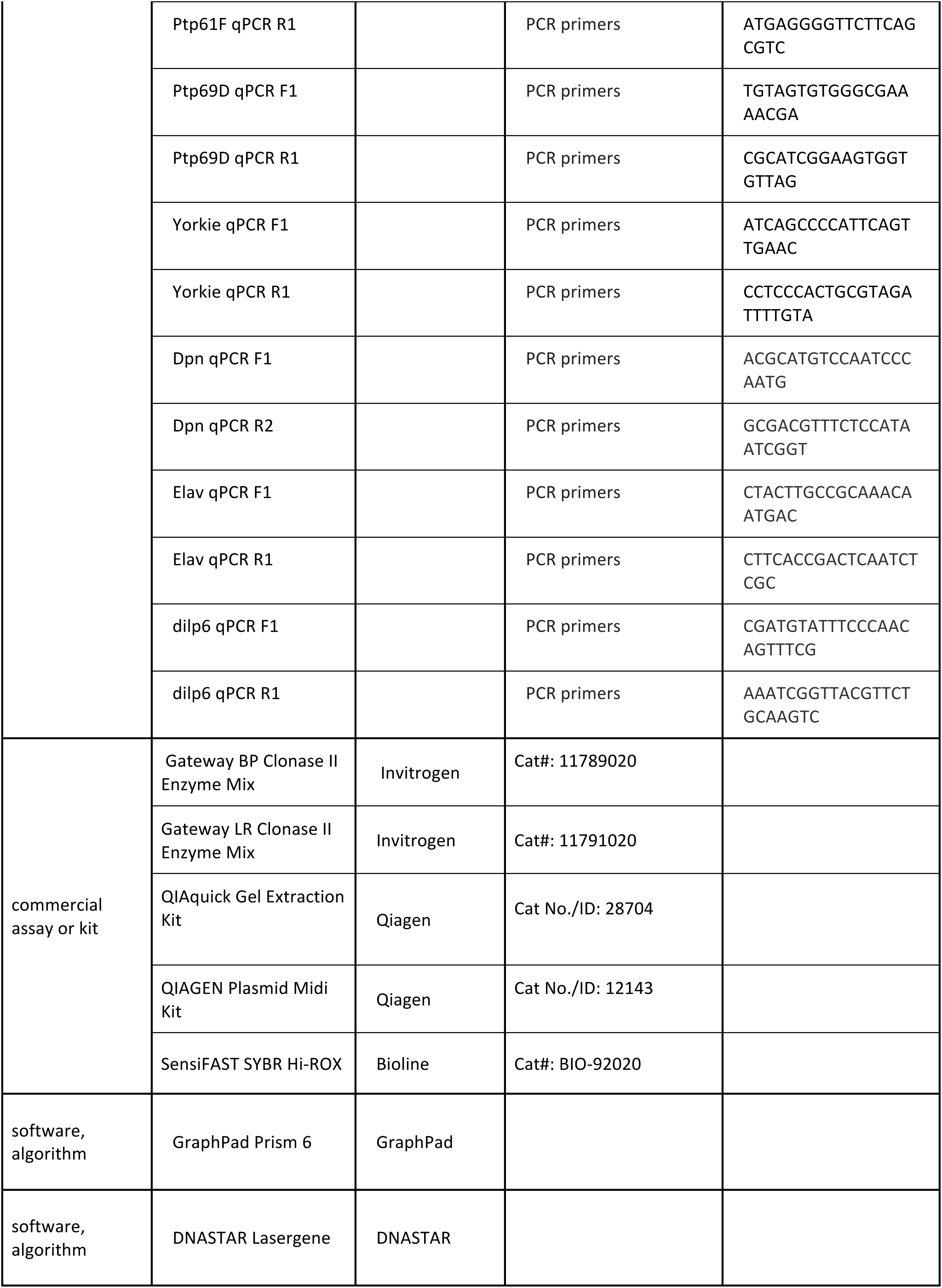
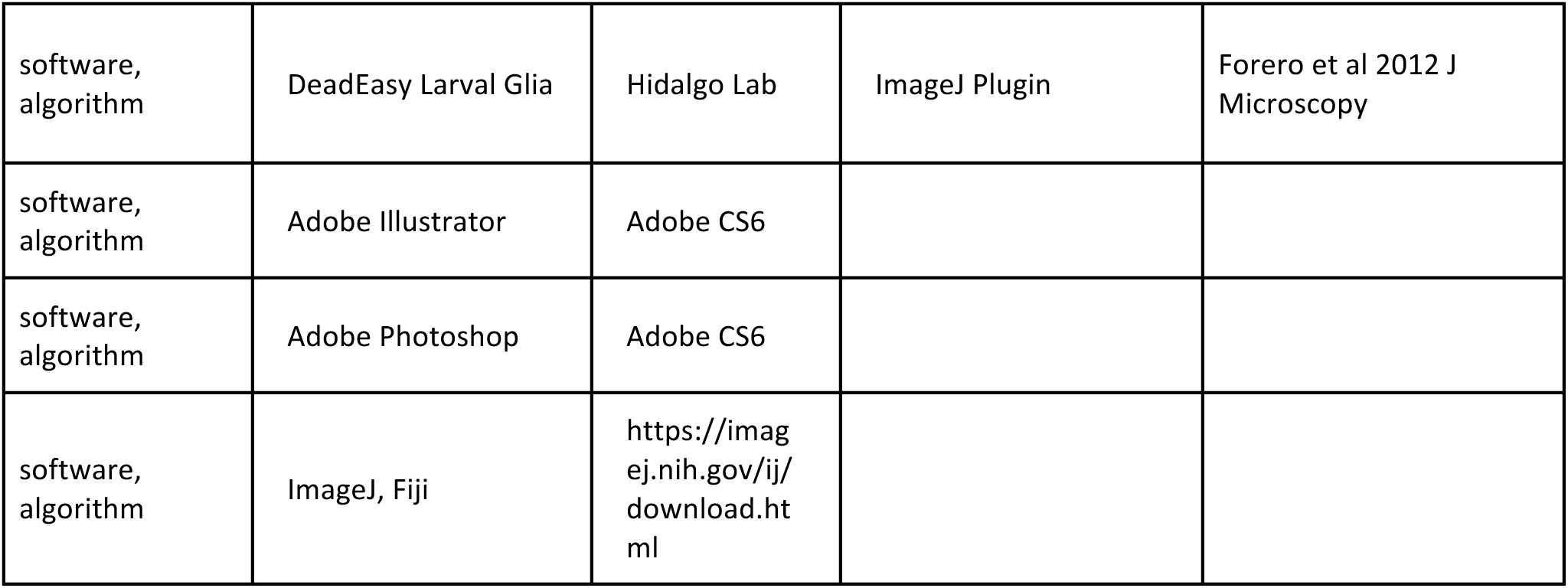
Key Resources Table.

